# Emergence of D1.1 reassortant H5N1 avian influenza viruses in North America

**DOI:** 10.64898/2025.12.19.695329

**Authors:** Alvin Crespo-Bellido, Nídia S. Trovão, Wendy Puryear, Alexander Maksiaev, Jonathan E. Pekar, Guy Baele, Simon Dellicour, Martha I. Nelson

**Author notes:** **Corresponding authors:** Alvin Crespo-Bellido and Martha I. Nelson.

## Abstract

Since 2021, highly pathogenic avian influenza viruses (HPAIVs) belonging to H5N1 clade 2.3.4.4b have circulated widely in North American wild birds and repeatedly spilled over into mammals. In 2025, the first H5N1-associated deaths in humans were recorded in the Western hemisphere, raising questions about how the ongoing evolution of the virus in wild birds impacts spillover risk. Here, our analysis of 21,471 H5N1 genomes identified an evolutionary shift in mid-2024, driven by interhemispheric migration from Asia and reassortment with new antigens. The genotypes that dominated the early years of North America’s H5N1 epizootic traced their ancestry back to Europe, but Asia was the source of new “D1.1” genotype viruses that (a) spread faster, (b) have higher reassortment potential, (c) a broader host range, (d) repeatedly spill over to bovines, and (e) cause severe disease in humans, including non-farm workers.

## Introduction

Highly pathogenic H5N1 avian influenza viruses from clade 2.3.4.4b have circulated in Eurasia since 2016^1^, causing severe mortality events in wild birds^2^, poultry^3,4^, and farmed mink and foxes^5,6^. In November 2021, H5N1 clade 2.3.4.4b viruses were introduced by wild birds migrating from Europe to Newfoundland, Canada, representing the first detection of clade 2.3.4.4b in the Western hemisphere^7^. The newly introduced H5N1 viruses spread across North America’s four migratory flyways during 2021-2022 and underwent genomic reassortment with North American low pathogenicity avian influenza (LPAI) viruses. To categorize the diversity of new H5N1 genotypes that acquired North American LPAI segments, a “GenoFlu” software tool was developed by researchers at the US Department of Agriculture to rapidly categorize new strains (https://github.com/USDA-VS/GenoFLU)^8^. Non-reassorted Eurasian H5N1 viruses are referred to as “A” genotypes. New reassortant genotypes generated by reassortment events between the A genotypes and LPAI are referred to as the “B, C, or D” genotypes. The B genotypes have been the source of most notable outbreaks in the Americas, including mass mortality events in South American coastal wildlife (B3.2)^9–12^, spillover to goats in Minnesota in March 2024 (B3.6)^13^, and a multistate outbreak in US dairy cattle that began in 2024 (B3.13)^14–16^. In 2024, a new D1.1 genotype emerged with a new N1 segment acquired from LPAI that has been highly costly for US poultry producers, caused the first human deaths by HPAI in the Americas, and spilled over multiple times into US dairy cattle^17^.

Whereas B3.13 spillover events primarily occurred in dairy and poultry workers and resulted in mild conjunctivitis symptoms, D1.1 zoonoses have been associated with more severe illness and the animal source has been less clear. The first D1.1 zoonotic infection involved a hospitalized teenager with no known animal exposure in British Columbia, Canada^18^ (November 9, 2024, **Figure S1**). In December 2024, an adult with underlying medical conditions died from a D1.1 infection following exposure to backyard poultry in Louisiana, USA, the first H5N1-associated fatality observed in the Americas^19^. In March 2025, a D1.1 infection in Durango, Mexico led to the death of a young child with no underlying medical conditions, no travel history, nor exposure to infected animals^20^. This death was the first reported H5N1 human infection in Mexico.

To determine how the ongoing evolution of H5N1 in North American wild birds impacts spillover risk for livestock, humans, and mammalian wildlife, we performed comparative and phylogenetic analyses using 21,471 H5Nx clade 2.3.4.4b genome sequences collected from all host species in the Americas, as of January 2, 2026. Bayesian phylodynamic approaches were used to infer evolutionary processes that drive H5N1 evolution in North America, including ongoing intercontinental migration, rapid spatial diffusion across the North American landmass, genomic reassortment with LPAI, and interspecies transmission involving host adaptation. In light of the sudden expansion of the D1.1 genotype in North America, we compared how D1.1 evolution differs from prior H5N1 genotypes across these four domains.

## Results

### Three introductions of H5Nx clade 2.3.4.4b viruses from Europe into North America’s eastern Atlantic flyway

Multiple independent introductions of clade 2.3.4.4b viruses from Eurasia into North America were identified by phylogenetic analyses of individual gene segments from 5,035 H5N1 viruses collected in Europe, Asia and North America (**Figure 1, Figure S2**). The first clade 2.3.4.4b viruses were introduced into North America’s Atlantic flyway from Europe in 2021 (genotype “A2”). A2 became the ancestor of the vast majority of H5N1 genetic diversity that evolved in North America during 2022-2025, including the numerous B and C genotypes that evolved via reassortment with North American LPAIs (**Figure 1, Figure S2**) that were dominant from early 2022 to mid-2024 (**Figure 2**). The C genotypes (e.g., C2.1) evolved independently from a genetically drifted descendent of A2 (referred to as “A1,” **Figure 1**). Genotypes A5 (H5N1) and A6 (H5N5) were independently introduced from Europe into North America’s Atlantic flyway but had no evidence of reassortment with North American LPAI (**Figures S2-S3**).

**Figure 1.**
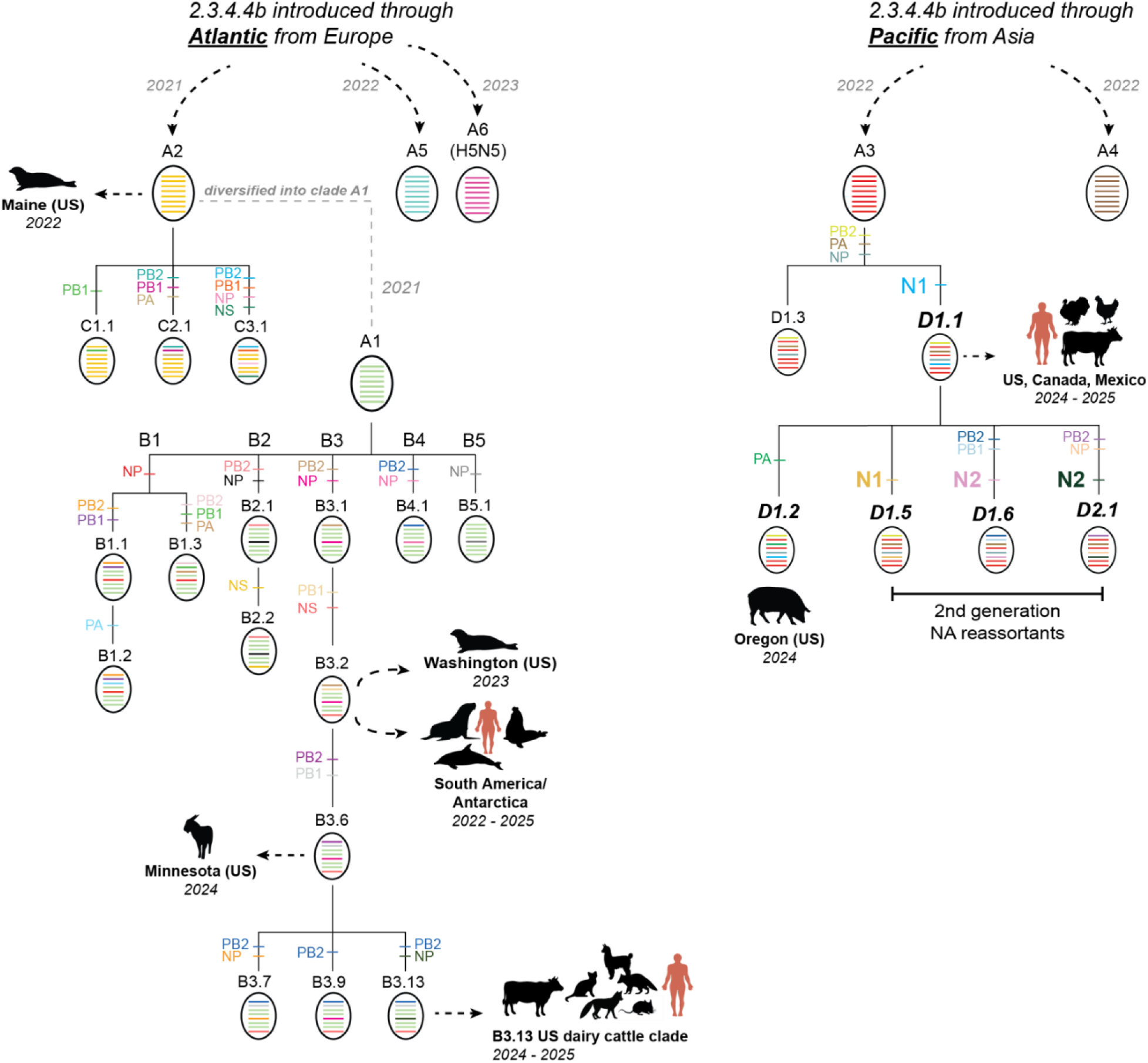
Reassortment history of H5 clade 2.3.4.4b genotypes in the Americas. Each oval represents a genotype (A-D). Each bar inside the oval represents one of the eight segments of the IAV genome, ordered from longest to shortest: PB2, PB1, PA, HA, NP, NA, MP, NS. Bars are shaded by lineage, as defined by GenoFlu (**Supplemental File 1**) and confirmed by phylogenetic analysis (**Figure S2**). For simplicity, all bars from Eurasian-origin “A” genotypes are shaded the same color. Horizontal bars with segment labels indicate where North American LPAI segments were acquired via reassortment, again colored by lineage. Dotted arrows and animal cartoons are provided for select outbreaks associated with spillovers to mammals.

**Figure 2.**
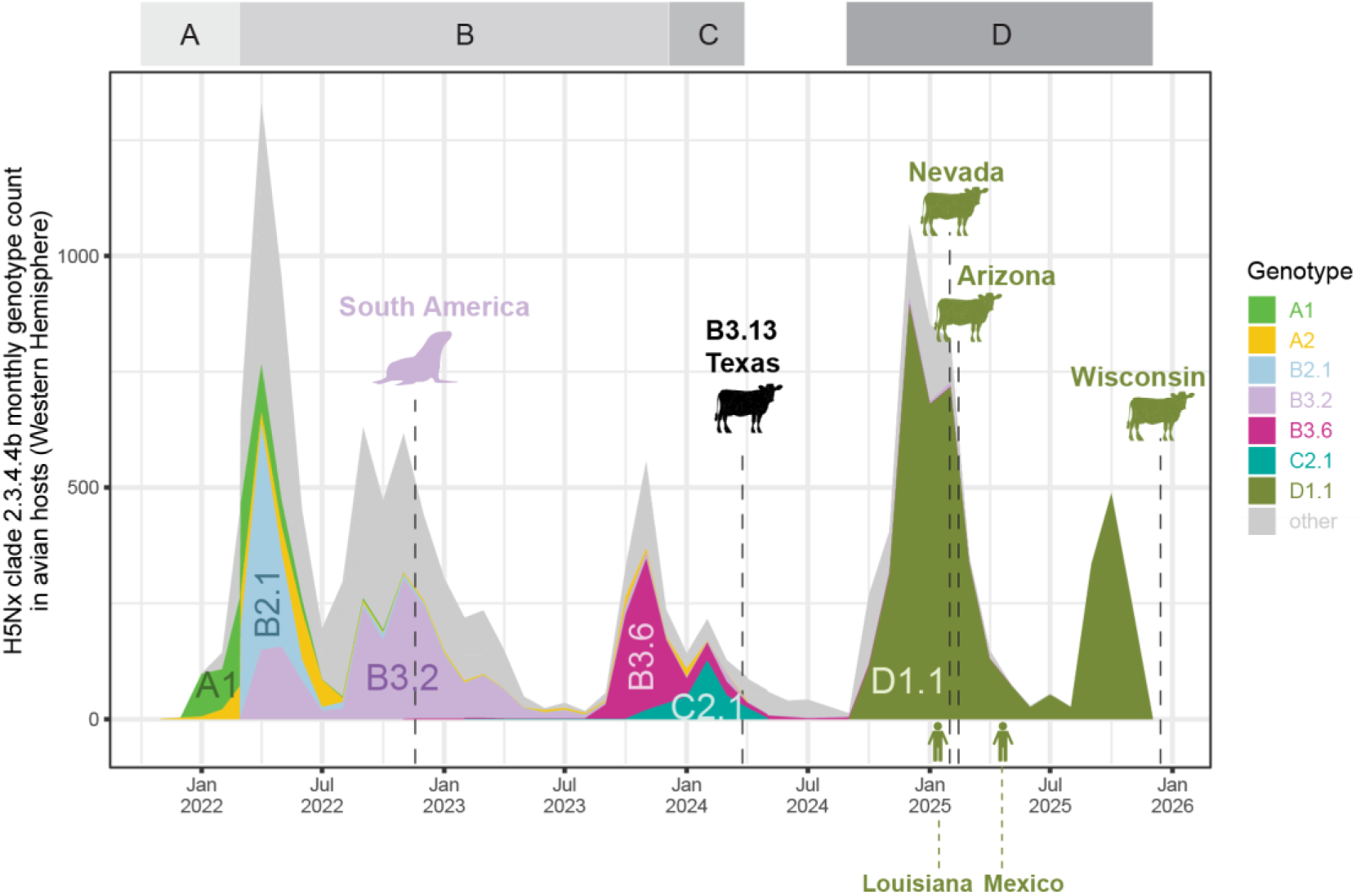
Timing of H5N1 reassortant genotypes in the Americas. The y-axis indicates the number of H5Nx clade 2.3.4.4b sequences detected for each genotype, by month, in avian hosts in the Americas during November 2021 - February 2025, based on 14,390 H5Nx 2.3.4.4b genomes with complete collection dates. The first detected virus is labeled with animal cartoons and dashed lines for select outbreaks/cases: B3.2 in South American marine mammals; B3.13 in Texas dairy cattle; D1.1 in dairy cattle in Nevada, Arizona and Wisconsin; and D1.1-associated deaths in Louisiana and Mexico. Time periods where A-D genotypes are dominant are labeled by boxes above the plot.

### Seven introductions of H5N1 clade 2.3.4.4b viruses from Asia into North America

Phylogenetic analysis reveals that the “A3” genotype was introduced from Asia into the Pacific flyway of North America (**Figure S4A**) at least six times independently during 2021-2024 (**Figure 3**). Five of the six A3 introductions (introductions 1, 2, and 4-6) did not sustain long-term transmission in North American wild birds or reassort with LPAI. The third A3 introduction, however, circulated widely in wild birds in the Pacific flyway, before undergoing genomic reassortment with North American LPAI to generate genotype D (**Figure 1**). The D1.1 genotype dominated the North American H5N1 epizootic during late 2024 and throughout 2025 and is the only genotype to dominate in North America during two successive fall migration seasons (2024 and 2025) (**Figure 2**).

**Figure 3.**
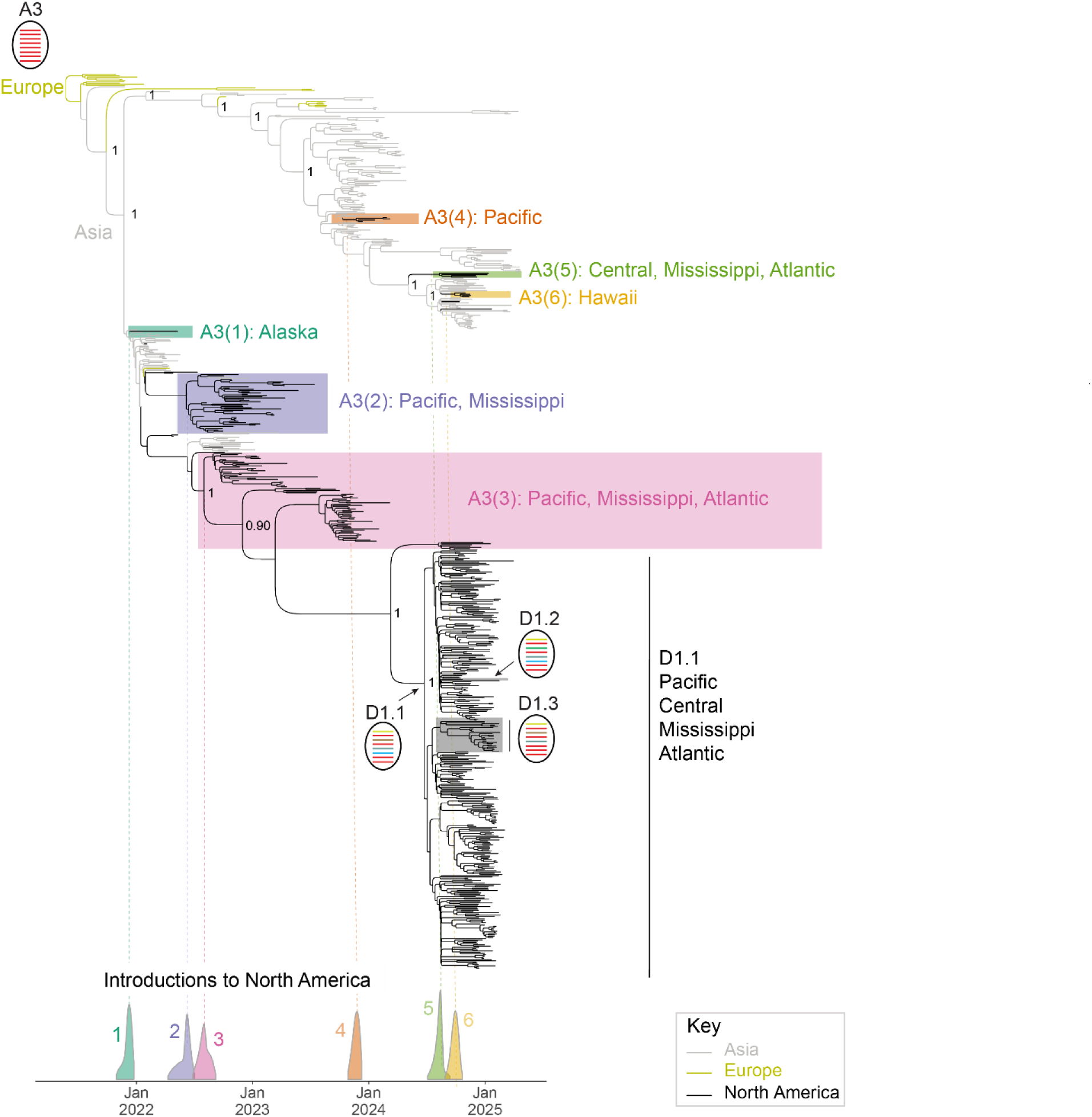
Introductions of A3 genotype into North American wild birds from Asia. Time-scaled maximum clade credibility (MCC) tree inferred for four concatenated genome segments (PB1, HA, MP and NS) for which the Eurasian lineage is retained in the D1.1, D1.2, and D1.3 genotypes. Branches are shaded by geographic location. Posterior probabilities are provided for key nodes. The 95% highest posterior density interval for the timing of each of the six A3 North American introductions is provided below the phylogeny. Schematics of reassortant genomes for the genotypes represented in the tree are similar to those presented in Figure 1.

### D1.1 acquires N1 from North American LPAI during reassortment

Prior to the emergence of D1.1 in October 2024, 100% of H5N1 viruses circulating in the Americas still retained the Eurasian N1 (**Figure S5**). Among clade 2.3.4.4b viruses sampled in North American wild birds in 2025 (*n* = 2,576), 92.4% contained an LPAI NA, dominated by D1.1 (**Figure S5**). Most reassortment events between H5N1 clade 2.3.4.4b viruses and North American LPAI during 2022-2025 involved the viral ribonucleoprotein (vRNP) complex, which includes the polymerase (PB2, PB1, PA) and NP segments (**Figure S4D-E**). NA and MP segments from LPAI were rarely acquired during reassortment. In a 4:4 reassortment event, the D1.1 genotype acquired the PB2, PA, NP, and N1 segments from LPAI (**Figure S4E**). D1.1 continued to reassort with LPAI to generate “second generation NA reassortants,” including D1.6 and D2.1 with N2 segments and D1.5 with yet another distinct N1. (**Figure 1**). The D1.2 virus that infected swine in Oregon in 2024 is similar to D1.2 but underwent another reassortment event in which a different PA was acquired.

### Mutations in D1.1 neuraminidase

The acquisition of an entire N1 segment from North American LPAI introduced a major genetic change into the H5N1 population. The LPAI N1 acquired by D1.1 viruses differs at 22 amino acid positions (**Table S2**) compared to the Eurasian-origin N1 segment found in other H5N1 genotypes in the Americas and globally (mean pairwise genetic percent similarity = 85.6%, **Figure 4A**). The 22 residues are distributed across all major functional domains of the N1 protein (**Table S3**), including 11 in the transmembrane and stalk domains (positions 1-84 in N1 numbering) that anchor the protein in the viral envelope and influence its stability, and 11 on the globular head domain (positions 85+ in N1 numbering) that is a primary target for the host immune system. The mutations on D1.1’s globular head domain are arranged in a diagonal pattern on the underside surface of the head, which typically is highly conserved (**Figure 4B**). How these NA mutations affect pandemic risk and immune recognition by antibodies elicited by humans against prior seasonal strains is an important question. The 84.1% genetic similarity observed between the D1.1 N1 and the 2009 H1N1 pandemic (H1N1pdm09) clade is similar to the 84.2% similarity between the B3.13 N1 and H1N1pdm09.

**Figure 4.**
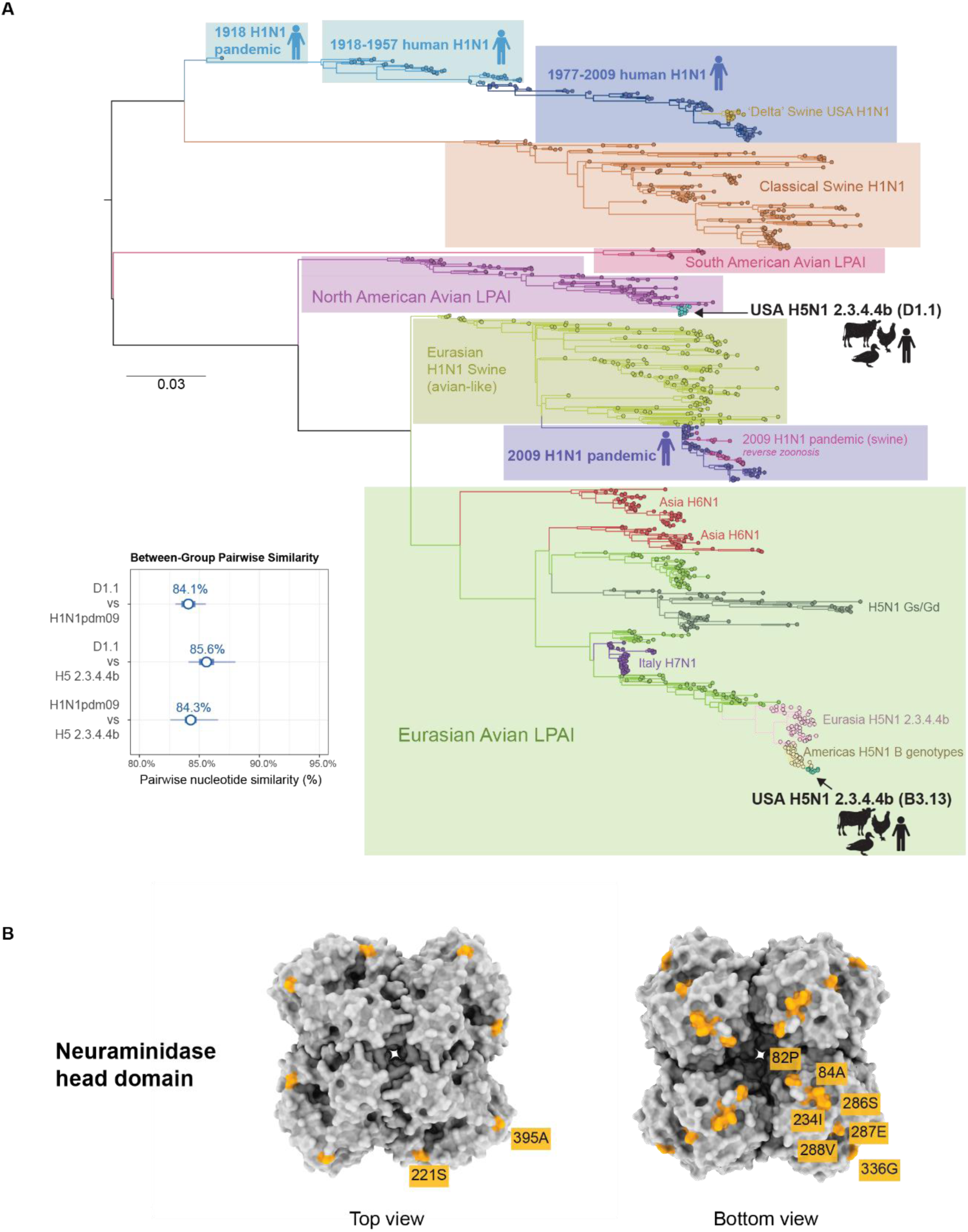
North American N1 segment acquired by D1.1 genotype. (A) Global phylogeny for N1 segment. Major N1 lineages as well as the placement of genotypes D1.1 and B3.13 are highlighted. The inset dot-range plot displays the mean pairwise similarity (dot), the mean ± one standard deviation (thick bar), and the full range of observed pairwise similarities (thin bar) for each between-group comparison. (B) Tetrameric NA head domain protein structure corresponding to H1N1 strain A/California/04/2009 (PDB: 3NSS) annotated with D1.1-specific residues. The surface of the head protein structure is shown at the top while the underside is shown at the bottom. Residues are listed according to N1 (sequential) numbering; the equivalent N2 numbering is listed in Table S3.

### Rapid spatial diffusion of D1.1 in North America

Our phylogeographic analysis inferred that the D1.1 genotype emerged in the Central flyway, or possibly Pacific flyway, during the summer of 2024 (95% HPD = 2024-06-17 to 2024-07-24; **Figures S6-S8**). The discrete trait analysis (DTA) provided higher support for the origin of D1.1 in the Central flyway (posterior probability of 0.70, **Figure 5A**), while a post-hoc test based on a continuous phylogeographic analysis provided moderate support for the Pacific flyway (posterior probability = 0.54) and Central flyway (posterior probability = 0.44), with uncertainty reflected in the 95% HPD polygon of the D1.1 MRCA in **Figure 6**. A demographic reconstruction of D1.1 indicates that effective population size increased sharply in the fall of 2024, declined during the spring/summer of 2025, and once again increased during the fall of 2025, consistent with the timing of fall migration (**Figure 5B**). D1.1 lineages tended to remain within a flyway more than expected by chance^21^, as evidenced by positive (but not strong) statistical support for a flyway containment effect (Bayes factor = 4.2). D1.1 appeared to persist in the Atlantic flyway during the summer of 2025 (**Figure 5A**). However, viruses sequenced during the second D1.1 wave primarily descended from viruses in the Central flyway positioned deeper in the tree. The continuous phylogeographic reconstruction estimated that D1.1 dispersed across all four flyways during 2024-2025 with the highest weighted diffusion coefficient of North America’s four major dominant genotypes during its invasion phase (∼6,000 km²/day, **Figure 6**, **Table S4**). The wavefront velocity during D1.1’s invasion phase (∼16,000 km/year) was substantially higher than the wavefront velocities for B3.2 (∼7,000 km/year), B3.6 (∼2,500 km/year) and C2.1 (∼6,000 km/year, **Figure 6**).

**Figure 5.**
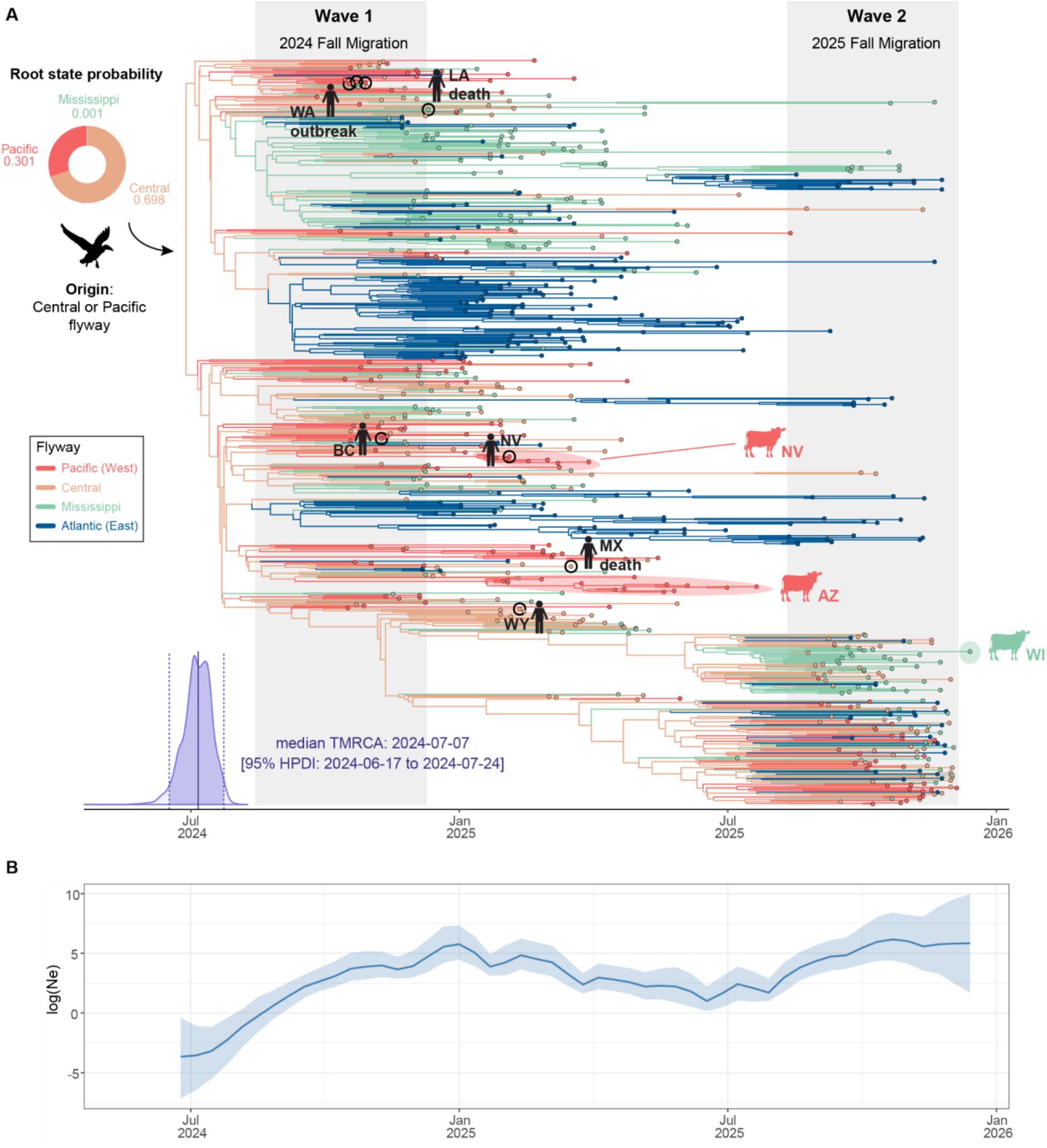
MCC tree of D1.1. (A) Discrete phylogeographic reconstruction of the dispersal history for H5N1 genotype D1.1 across the North American waterfowl migratory flyways. The consensus phylogenetic tree shows the evolutionary relationships of genotype D1.1 viruses, with the x-axis scaled in years. Tips are shaded according to the flyway where they were collected, and branches are shaded according to the flyway inferred at ancestral nodes. The purple curve at the root of the tree represents the posterior density distribution for the TMRCA. (B) Skygrid reconstruction of the effective population size for D1.1. The dark blue line and shaded light blue region represent the median log-transformed viral effective population size *Ne* and its 95% highest posterior density (HPD) interval region, respectively.

**Figure 6.**
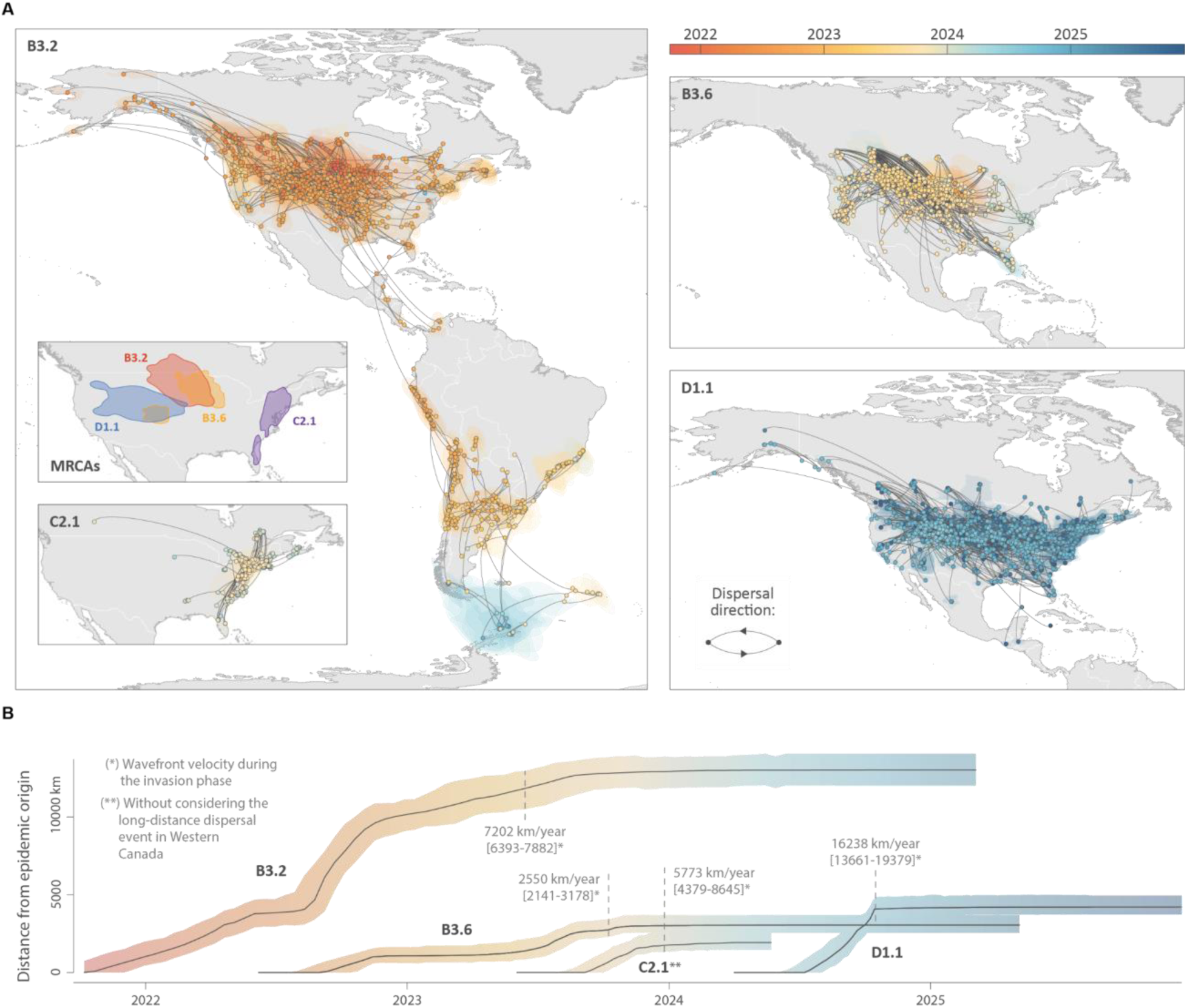
Spatial dispersal of major H5N1 genotypes in the Americas. (A) Phylogeographic reconstruction maps showing maximum clade credibility (MCC) tree branches as curved lines. Nodes are time-colored from red (2022) to blue (2025), with 80% highest posterior density (HPD) polygons indicating uncertainty. Inset maps show the estimated geographic origins as the 95% HPD regions for most recent common ancestors (top) and the C2.1 phylogeographic reconstruction (bottom). (B) Expansion dynamics showing median wavefront distance from geographic origin over time, with 95% HPD intervals. The annotated values indicate median wavefront velocities (km/year) during the initial invasion phase.

### D1.1 spillover and adaptation to mammals

D1.1 has frequently transmitted from wild birds to a range of diverse host species (**Figure 7A, Figure S9**). D1.1 has a broad host range compared to prior genotypes and is the only genotype to infect seven host types: wild bird, terrestrial mammal, marine mammal, domestic bird, cattle, domestic mammal (other than cattle), and human (**Figure 7B**). D1.1 also infected a broader range of wild bird orders (*n* = 14, **Figure 7C**) than any other genotype, including the B3.2 genotype that spread widely in South America. Ecologically, most D1.1 spillovers involved direct exposure to wild birds (**Figure 7A**) and these avian-origin viruses experienced strong selection in mammalian hosts for PB2 mutations that improve virus replication efficiency in mammals. Close to 35% of D1.1 spillovers from wild bird into mammals (e.g., humans, seals, bears, raccoons, skunks) had the E627K mutation in PB2, including the human cases A/Wyoming/01/2025 and A/British Columbia/PHL-2032/2024 (**Figure 8, Figure S10**). The D701N mutation in PB2 seen in South American marine mammals also became fixed in the D1.1 Nevada cattle clade and was found in ∼7% of other D1.1 spillovers into mammals (e.g., domestic cat, fox). In contrast, once the PB2 segment became adapted to cattle (D701N for D1.1; M631L for B3.13), there is no evidence of emergence for additional PB2 mutations following spillover events from cattle to other mammalian hosts (**Figure 8, FiguresS10-S11**).

**Figure 7.**
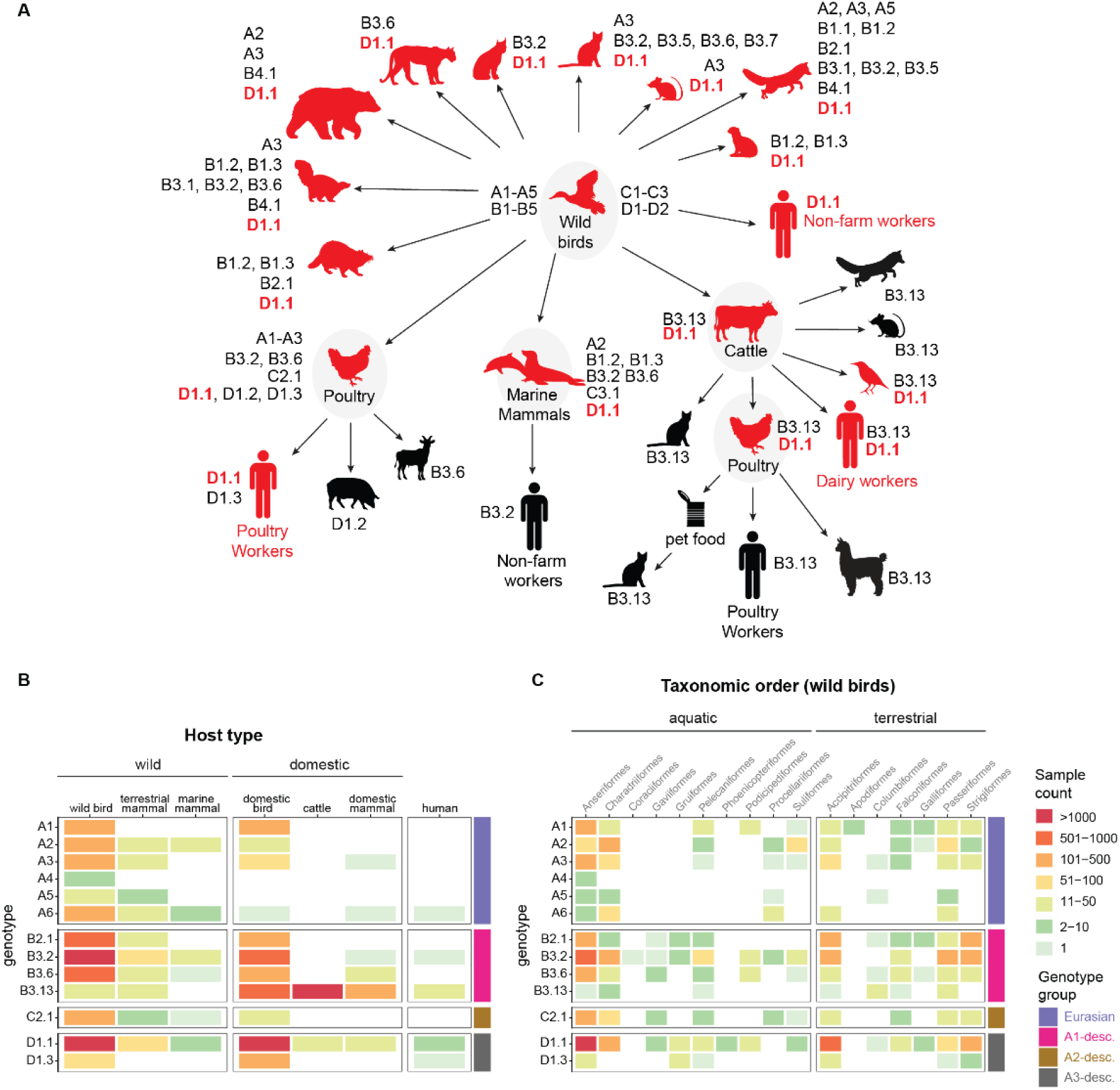
Host range of H5N1 clade 2.3.4.4b genotypes. (A) Spillover of H5N1 clade 2.3.4.4b genotypes between different host species based on phylogenetic evidence. (B) Heatmap shaded by the number of sequences in each wild bird taxonomic order (*n* = 4,044) and (C) host type (*n* = 13,488). Select genotypes are shown.

**Figure 8.**
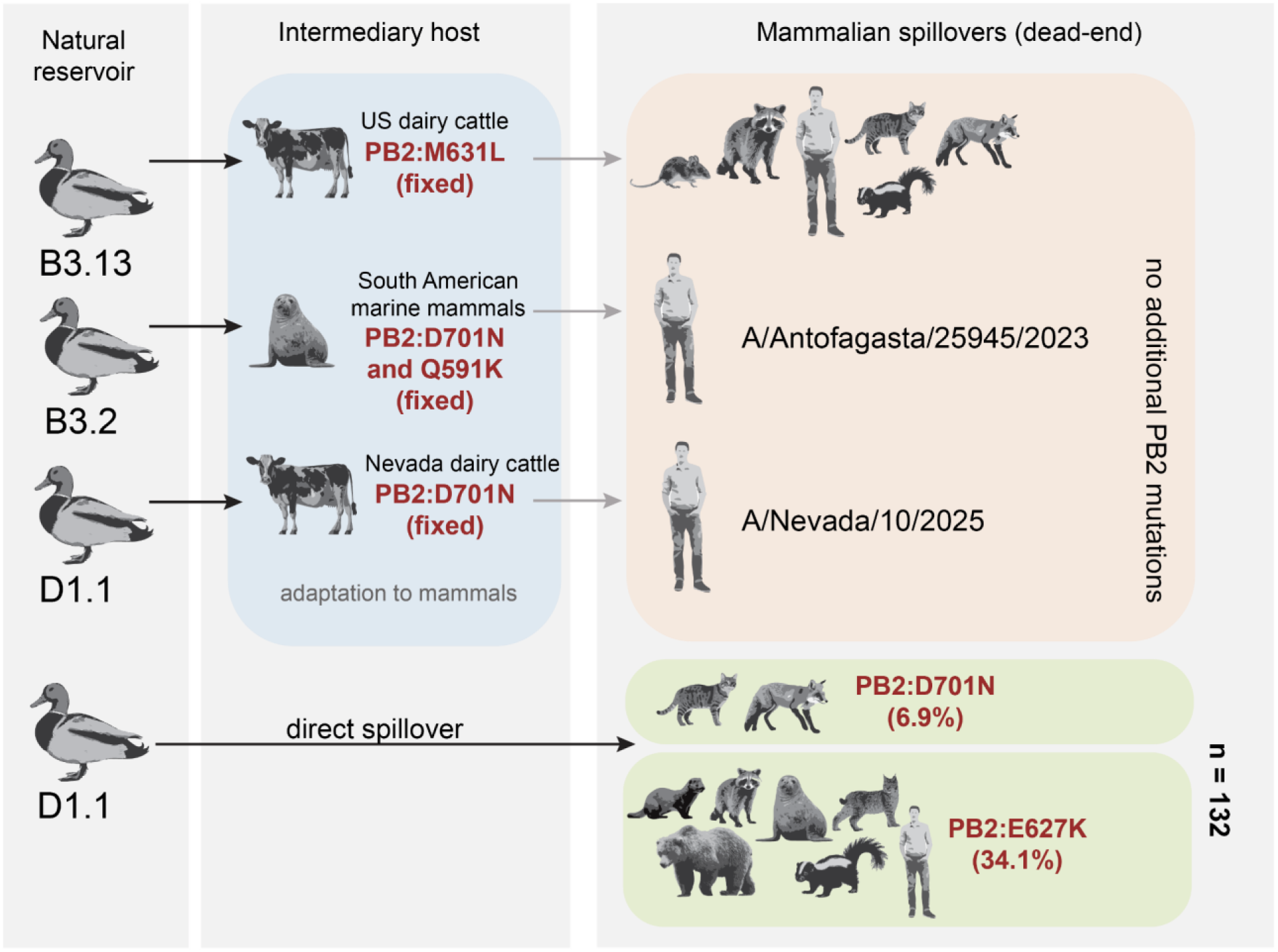
PB2 mutations following spillover events in mammals. Black arrows indicate H5N1 spillover events from the avian natural reservoir to mammalian hosts. Grey arrows = secondary spillover events from intermediary hosts (e.g., bovines, marine mammals) to dead-end mammalian hosts. Blue box = intermediary hosts that acquired PB2 mutations that improve virus replication in mammals. Orange box = secondary hosts that experience little to no selection for additional PB2 mutations because viruses transmitted from intermediaries are already adapted to mammals. Green boxes = mammals infected with avian-origin D1.1 where selection is strong for PB2 adaptations.

## Discussion

The emergence of the D1.1 genotype reflects North America’s expanding role as a source for novel reassortant H5N1 viruses with pandemic potential and the increasing complexity of the H5N1 epizootic’s geography, genetics, and spillover risk. Our findings support a model in which global viral connectivity introduces genetically distinct lineages into new ecological systems, where reassortment with local gene pools generates genotypes with enhanced evolutionary flexibility, enabling rapid expansion and cross-species transmission. Prior to the emergence of D1.1 in 2024, the genetic diversity of the North American 2.3.4.4b epizootic was constrained by the fact that (a) almost all strains traced their ancestry back to the original 2.3.4.4b introduction from Europe and (b) LPAI internal gene segments were almost exclusively acquired during reassortment events, with no swapping of NA antigens^22^. The human population at risk for spillover was likewise constrained, as H5N1 zoonotic cases in 2024 primarily occurred in US farm workers with direct animal contact and generally caused mild illness. It remains unclear whether the H5N1 fatalities caused by D1.1 are an indication of a higher pathogenicity of the virus in humans or a larger denominator of human cases beyond just farm workers. In which case H5N1 surveillance in humans has not kept pace with the expanding population at risk.

D1.1 is the only genotype to cause two successive epidemic peaks in wild birds, with no sign of being replaced by a different genotype. Prior North American genotypes dominated for one epidemic peak (although B3.2 continues to circulate in South America and the Antarctic region^23^). At the time of this writing, D1.1 is continuing to acquire new N1 and N2 antigens through additional reassortment events with LPAI, suggesting that H5N1 evolution is continuing to accelerate and explore new antigenic pairings.

Following years of high conservation of the HA-NA pairing of 2.3.4.4b viruses, it is unclear why a new North American LPAI N1 was acquired in 2024 without any apparent loss in fitness. We found no evidence of permissive mutations^24^ in the Asian-origin A3 hemagglutinin in the months leading up to D1.1’s emergence that could have explained a greater tolerance for a new NA. The implications of the new N1 for pandemic risk require further research. Human exposure to H1N1 seasonal influenza viruses containing N1 segments from the 2009 H1N1 pandemic, originally sourced from Eurasian swine, is thought to provide a degree of cross-protection for humans against most H5N1 clade 2.3.4.4b viruses circulating in birds globally. The amino acid changes that distinguish the LPAI N1 found in D1.1 from the Eurasian N1 are concentrated on the underside of the globular head domain – or the neuraminidase “dark side” (NDS) – which is positioned away from the enzymatic active site. This region has recently been identified as containing highly conserved epitopes that serve as effective targets for cross-reactive monoclonal antibodies against HxN1 infections^25^.

As the Americas emerge as a new epicenter for H5N1 evolution, there is an urgent need to expand the region’s infrastructure for real-time surveillance and monitoring. The Americas present a different ecological landscape compared to Eurasia, including South American marine mammals that are highly susceptible to H5N1 disease and farming systems with long-range animal movements that sustain virus persistence in dairy cattle^16^. Tracking H5N1 evolution in the Americas is made possible by the collective efforts of government and academic partners, including the U.S. Department of Agriculture^8^, Centers for Disease Control and Prevention^26^, US Geological Survey^27^, the Canadian government^28^, South American researchers^10,12^, and NIAID-funded Centers of Excellence for Influenza Research and Response (CEIRR) network^29^. The capacity of the USDA’s National Milk Testing Strategy (NMTS) to detect three independent D1.1 spillover events into US dairy cattle by actively sampling raw bulk milk tanks demonstrates one area where H5N1 active surveillance in North America is advancing. Still, there are severe gaps. A human fatality involving a highly pathogenic H5N5 virus in Washington State (genotype A6) was reported on November 15, 2025 (https://doh.wa.gov/newsroom/grays-harbor-county-resident-dies-complications-avian-influenza), despite no detections of H5N5 at that time in any host outside of the Atlantic flyway on the opposite coast^30^, exposing major gaps in our knowledge of which genotypes are circulating in wild birds (and where) caused by a lag in H5N1 data submission to public repositories. Numerous genotypes are listed on GenoFlu for which no or very few genetic sequences are available yet on SRA or GenBank, including some of the latest D genotypes with novel NA segments that could not be included in this study. Tracking how genotypes evolve over time and space is also limited by the large numbers of H5N1 genomes on SRA that are labeled “USA” and have no collection date. It can take months (or more) before a corresponding GenBank entry appears with state and date metadata. H5N1 surveillance is severely underfunded, but it would be sensible to invest more resources into high-throughput data pipelines so that crucial H5N1 data already being generated by government agencies can quickly get into the hands of the global scientific community to track changes in evolution and spillover risk in real time.

In recent years, North American avian influenza surveillance has prioritized sequencing HPAI over LPAI, for good reason. However, LPAIs still present an important pandemic risk (e.g., 1918, 1957, 1968, and 2009 influenza pandemics) and contribute key segments to new H5N1 reassortants and should not be overlooked. Lower availability of LPAI sequences from countries around the world in public databases decreases phylogenetic resolution and makes it difficult to distinguish reassortment events between HPAI and LPAI. One source of confusion is the A1 and A2 genotypes and whether they represent the first two 2.3.4.4b introductions from Europe. Our updated phylogenetic analysis supports an alternate scenario where A1 and A2 descend from the same introduction from Europe. A2 arrived first, and A1 genetically drifted into a new lineage as mutations accumulated over time (**Figures 1 and S2**). Efforts to develop a new universal global H5N1 lineage classification tool (“ggFlu”, https://www.ggflu.com/) represent the latest attempt to bring order to the increasing complexity of global H5N1 evolution (**Table S5**).

## Methods

### Dataset curation and maximum likelihood phylogenetic analysis for H5 clade 2.3.4.4b

An H5Nx clade 2.3.4.4b dataset for North America, South America and Antarctica was compiled by querying the Global Initiative on Sharing All Influenza Data (GISAID; https://gisaid.org/)^31–33^ and NCBI Virus^34^ databases for all H5Nx clade 2.3.4.4b samples collected between November 1, 2021 – January 2, 2026 with available genomic sequences for all eight gene segments (accessed on June 2, 2026). Additionally, consensus genomes from raw reads deposited to NCBI’s Sequence Read Archive and associated with BioProjects PRJNA1102327, PRJNA1122849, PRJNA1134696, PRJNA1219588, PRJNA1207547, PRJNA980729 were downloaded from the Andersen Lab’s avian-influenza github repository (https://github.com/andersen-lab/avian-influenza). The samples in each dataset were assigned a genotype using the GenoFLU genotyping tool^8^. GenoFLU categorizes reassortant clade 2.3.4.4b genotypes as “major” (*n* = 37, as of April 7, 2026) if they meet the criteria of at least 20 wild bird detections, infection of two or more poultry premises, and/or detection in atypical mammalian host species. The vast majority of reassortant genotypes (123/160) are designated by GenoFlu as ‘minor’. Samples with unassigned genotypes were removed from the dataset and duplicate samples across databases were filtered by isolate id. Of the 166 genotypes listed on GenoFLU’s website, genetic sequence data was publicly available for 107 of them (∼65%). The final Western Hemisphere clade 2.3.4.4b dataset consisted of 21,471 genome sequences. To place the clade 2.3.4.4b genotypes in phylogenetic context and identify reassortment involving American LPAI viruses, a background dataset was compiled by querying the GISAID database for LPAI samples with complete genomes collected in the Americas and Antarctica between January 1, 2015 – January 2, 2026 (accessed on April 4, 2026). Select samples belonging to distinct lineages were removed from the analysis which included H1N1 and H1N2 samples from swine, and H3N8 samples from canines and horses. The search resulted in an LPAI dataset of 3,830 samples. All sequences from the American clade 2.3.4.4b and background LPAI were combined and grouped by segment resulting in eight segment datasets – PB2, PB1, PA, H5, NP, N1, MP and NS.

An additional background dataset was compiled to evaluate the number of North American clade 2.3.4.4b introductions of fully Eurasian genotypes A1-A6 by downloading all European and Asian H5Nx clade 2.3.4.4b and LPAI samples with complete genomes collected between January 1, 2015 and June 13, 2025 (accessed on June 13, 2025). To reduce the size of the datasets while keeping a representative Eurasian background sample set, the sequences were then grouped by segment and used to create local Basic Local Alignment Search Tool (BLAST) databases using BLAST v2.15.0^35^. The resulting Eurasian segment databases were BLASTed against using the American A1-A6 clade 2.3.4.4b datasets as queries, and the top 200 Eurasian BLAST hits with >85% sequence identity across all American clade 2.3.4.4b sequences were included for subsequent phylogenetic analysis. The final American A1-A6 + Eurasian sequence datasets consisted of: H5 (*n =* 4,923), MP (*n =* 5,035), N1 (*n =* 3,935), NP (*n =* 5,035), NS (*n =* 5,035), PA (*n =* 5,035), PB1 (*n =* 5,035), and PB2 (*n =* 5,035).

Each gene segment dataset was aligned using MAFFT^36,37^ and trimmed to the coding sequence region. Highly diverged sequences that were poorly aligned were removed from each segment dataset. Sequences were also excluded excluded if more than 25% of bases were ambiguous or if their ungapped length fell below 75% of the mean ungapped length for that segment. Maximum-likelihood (ML) phylogenies were inferred for each dataset using IQ-TREE v2.47^38^ with a general-time reversible (GTR) model of nucleotide substitution with among-site rate heterogeneity modelled through a discretized gamma distribution (+G) and 1,000 ultra-fast bootstrap (UFBOOT) replicates, using the high-performance computational capabilities of the Biowulf Linux cluster at the National Institutes of Health (http://biowulf.nih.gov).

To increase the readability of the American clade 2.3.4.4b and background LPAI phylogenies consisting of >21,000 sequences, ML phylogenetic analyses were also performed on representative, subsampled datasets. Each segment dataset was subsampled using SAMPI (https://github.com/jlcherry/SAMPI)^39^. Subsampling was stratified by collection date, location, host category (avian, cattle, environmental, feline, human, and other mammals), and assigned genotype to maintain representative diversity across these variables. The resulting subsampled datasets consisted of H5 (*n =* 1,160), MP (*n =* 1,340), N1 (*n =*1,120), NP (*n =* 1,354), NS (*n =* 1,346), PA (*n =* 1,358), PB1 (*n =* 1,353), and PB2 (*n =* 1,372). Phylogenies were visualized using the “ggtree” R package v3.13.1^40^.

### Bayesian phylodynamic analyses

Bayesian phylodynamic reconstructions were performed for temporally dominant genotypes B3.2, B3.6, C2.1 and D1.1 using concatenated genome datasets (all eight segments were concatenated using ConcatenatorGUI^41^). For computational efficiency, subsampled sequence datasets were first obtained by conducting subsampling by phylogenetic clustering. This subsampling procedure consists in discarding sequences such that monophyletic clusters of sequences sampled from the same location (i.e. the most precise administrative region of origin of each sample) are only represented by a single sequence^42^. Given that such clusters would largely represent dispersal within a specific administrative region and given that most geographic coordinates are here integrated through a homogeneous prior range delineated by the polygon of this administrative area (see below), conserving more than sequence per cluster would lead to phylogeographic ‘noise’ and not further inform the overall dispersal history of viral lineages. For genotypes B3.2 and D1.1, we further decreased the size of the resulting datasets by randomly subsampling two sequences per combination of sampling month and region of administrative level 1 (admin-1 level), while preferentially conserving samples associated with the most precise sampling origins. This resulted in the following final datasets – B3.2 (*n* = 594), B3.6 (*n* = 526), C2.1 (*n* = 165), D1.1 (*n* = 699). To assess the timing of North American A3 introductions and the emergence of the D1 genotypes, an additional analysis was performed on a concatenated shared segment dataset (i.e., PB1, HA, MP and NS) of all Eurasian (downloaded from GISAID) and North American A3 sequences and a random subset of the D1.1-D1.3 samples (*n* = 725).

Bayesian phylogenetic and phylogeographic analyses were performed using the software package BEAST X v1.10.5^43^, using the BEAGLE high-performance library for computational efficiency^44^ and employing default prior settings in BEAUti X^43^ for all analyses. We employed a GTR+Γ nucleotide substitution model, an uncorrelated relaxed molecular clock with an underlying lognormal distribution to account for rate variation across lineages^45^, and a non-parametric SkyGrid coalescent tree prior^46^. Markov chain Monte Carlo (MCMC) chains were run for at least 300 million iterations while sampling every 10^5^ iterations. Empirical tree distributions of 1,000 trees were collected and subsequently used in the phylogeographic reconstructions^47^. To reconstruct the finer-scale geographic spread of the more recently emerged genotypes B3.2, B3.6, C2.1, and D1.1, we conducted continuous phylogeographic analyses using the relaxed random walk (RRW) diffusion model implemented in the software package BEAST X^43^, with a Cauchy distribution to model dispersal velocity heterogeneity^48–50^. For the genomic samples that did not have associated precise sampling coordinates, we used the polygon(s) of their administrative area of origin to define an homogeneous prior range of sampling coordinates and estimate those sampling locations through Bayesian phylogeographic inference^51,52^. We also conducted discrete phylogeographic analyses using the Bayesian stochastic search variable selection (BSSVS) approach^53^ implemented in BEAST X and while considering as a distinct discrete location each of the four administrative waterfowl flyways (i.e. the Pacific, Central, Mississippi and Atlantic flyways, as defined by the U.S. Fish and Wildlife Service). For all phylogeographic analyses based on empirical trees, MCMC chains were run for more than 10^7^ iterations while sampling every 10^5^ iterations. MCMC convergence and mixing were evaluated in Tracer v.1.7.3^54^, ensuring that all continuous posterior estimates were associated with an effective sample size (ESS) value of at least 200. At least 10% burn-in was removed from each run and maximum clade credibility (MCC) trees were computed using TreeAnnotator X^55^. We used the R package “seraphim”^56^ to generate a visualization for the different continuous phylogeographic reconstructions and for estimating dispersal statistics based on these reconstructions. For C2.1, a unique long-distance dispersal event detected towards Western Canada was removed from the dispersal calculations.

### Global N1 phylogeny and Pairwise nucleotide percent similarity between major N1 lineages

Similarly to Trovão et. al^57^, a background of all available neuraminidase sequences for influenza A N1 from GenBank and downloaded specific geographical isolates, not available in GenBank, from GISAID were obtained on 16 July 2019. The sequences were aligned using the MAFFT^36,37^ multiple sequence alignment tool. The aligned sequences were manually edited and cleaned in AliView version 1.26 software^58^. The resulting data set was trimmed at the 5′ and 3′ ends to include solely the coding sequence. This data set was subjected to multiple iterations of phylogeny reconstruction using FastTree version 1.0^59^ with a general time-reversible (GTR) model and exclusion of outlier sequences whose genetic divergence and sampling date were incongruent using TempEst^60^.

We defined lineages from the same host and geographical region and only considered those circulating for at least 3 years in the region. Each lineage includes only viruses that infect a single host species or closely related host species. Each inferred introduction into a new species creates a new lineage. In some cases, what could be a lineage is subdivided on the basis of geography. Besides the H1N1 pandemic lineage circulating in humans since 2009, we defined two human N1 lineages circulating during the periods of 1918-1957 and 1977-2009. The N1 lineage reintroduced in 1977 derives directly from the viruses circulating in 1957, so studying the two lineages together would disrupt the temporal signal and consequently the molecular clock model, due to the lack of accumulation of mutations during the 20-year period where N1 did not circulate^61–64^. For the non-avian lineages, we excluded sequences with mixed subtypes, as marked by “Nx” or “mixed.” For some analysis, we combine lineages for the same host. For visualization purposes, we excluded sequences that cover less than 75% of the coding sequence with unambiguous bases and retained only one of identical sequences collected in the same country on the same day or in the same month or year in the case of incomplete date information.

Next, we applied a subsampling strategy that selected a specific number of sequences per country per time to obtain similarly sized lineages that represented the overall circulating diversity. When sequences had the same country and collection date, we chose, in a decreasing order of priority, those that were longer, those that had at least the month of collection rather than just the year, those with a smaller number of internal gaps with lengths not divisible by 3 (codons), and those with full dates of collection (https://github.com/jlcherry/SAMPI)^39^. We combined all subsampled lineages for each subtype and performed a final iteration of phylogeny reconstruction and exclusion of outlier sequences as described above.

The abovementioned previously curated dataset was used to place the D1.1 and other clade 2.3.4.4b sequences in phylogenetic context. Pairwise nucleotide sequence similarity was calculated between all inter-group sequence pairs across the aligned N1 segment using a custom R script. For each group comparison, similarity was computed as the proportion of identical sites at non-gap, non-ambiguous positions across all pairwise combinations of sequences between the two groups.

### Identification of genotype-specific amino acid residues and D1.1 phenotypic markers

American H5Nx clade 2.3.4.4b segment datasets were translated into protein alignments with Nextclade CLI^65^ with built-in reference datasets: H5N8 ‘A/Astrakhan/3212/2020(H5N8)’ for HA, H1N1 ‘A/Wisconsin/588/2019’ for NA, and H1N1 ‘A/California/07/2009’ for all other segments. Sequences that failed to be aligned to the reference were not considered for this analysis. Genotype-specific amino acid residues were identified among dominant genotypes (B3.2, B3.6, B3.13, C2.1, and D1.1) using a custom R script. We defined genotype-specific residues as those meeting three criteria: (1) present in >90% of sequences within the target genotype, (2) present in <20% of other dominant genotypes, and (3) present in <10% of all other clade 2.3.4.4b sequences combined.

### Analysis of mammalian adaptation markers

Known phenotypic markers of mammalian adaptation (PB2:E627K and PB2:D701N) were identified within complete nucleotide alignments for D1.1 (n=5,257) and B3.13 (n=5,280) genotypes using FluMutGUI v3.2.0^66^. Maximum likelihood phylogenies were constructed from the D1.1 dataset using IQTREE as described above to characterize spillover events. Spillover events involving mammalian hosts were categorized by source as “direct spillover” from avian hosts, “AZ cattle clade”-derived, or “NV cattle clade”-derived. For B3.13, groups were defined as “pre-PB2:M631L” or “post-PB2:M631L”, given that M631L was fixed early in the cattle outbreak and subsequent spread was driven by spillovers from cattle^14^. We quantified the frequency of PB2 mutations across these spillover categories and different host types.

## Data and code availability

All data (including supplemental files and GISAID acknowledgement table) and BEAST X XML files with model parametrizations, phylogenetic trees and analysis output files are available at https://github.com/nidiatrovao/H5N1Americas.

## Acknowledgements

This research was supported in part by the Intramural Research Program of the National Institutes of Health (NIH). The contributions of the NIH author(s) are considered Works of the United States Government. The findings and conclusions presented in this paper are those of the author(s) and do not necessarily reflect the views of the NIH or the U.S. Department of Health and Human Services, or the United States government. We gratefully acknowledge all data contributors, i.e., the Authors and their Originating laboratories responsible for obtaining the specimens, and their Submitting laboratories for generating the genetic sequence and metadata and sharing via the GISAID Initiative, on which this research is based. We thank Dr. Joshua L. Cherry (NIH) for providing a curated global N1 dataset to place the clade 2.3.4.4b viruses in phylogenetic context. We also thank Dr. Andrew Ramey (USGS) and Dr. Christina Ahlstrom (USGS) for providing additional metadata on sequences collected by the USGS. GB and SD acknowledge support from the Research Foundation — Flanders (*Fonds voor Wetenschappelijk Onderzoek — Vlaanderen*, FWO, Belgium; grant n°G098321N), and from the European Union Horizon 2023 RIA project LEAPS (grant agreement no. 101094685). GB also acknowledges support from the Research Foundation — Flanders (*Fonds voor Wetenschappelijk Onderzoek — Vlaanderen*, FWO, Belgium; grant n° G0E1420N), and from the DURABLE EU4Health project 02/2023-01/2027, which is co-funded by the European Union (call EU4H-2021-PJ4) under Grant Agreement No. 101102733. SD also acknowledges support from the *Fonds National de la Recherche Scientifique* (F.R.S.-FNRS, Belgium; including grant n°F.4515.22), and from the University of Brussels (ULB) internal fund. This work is supported by the Centers of Excellence for Influenza Research and Response, National Institute of Allergy and Infectious Diseases, National Institutes of Health (NIH), Department of Health and Human Services, under contract 75N93021C00014.

## SUPPLEMENTARY MATERIAL

**Table S1.** List of H5Nx clade 2.3.4.4b with NA segments acquired through reassortment with North American LPAI. Genotype D1.1 and its major genotype descendants are in bold. GenoFLU-assigned lineages

| Genotype | Ancestry | Subtype | PB2 | PB1 | PA | HA | NP | NA | MP | NS | # of sequences |
| --- | --- | --- | --- | --- | --- | --- | --- | --- | --- | --- | --- |
| B3.14 | B3.2 descendant | H5N2 | am2.1 | mx1 | ea1 | ea1 | mx1 | mx1N2 | ea1 | am1.1 | 1 |
| **D1.1** | **D1.1 descendant** | **H5N1 (am-NA)** | **am24** | **ea3** | **am4** | **ea3** | **am13** | **am4N1** | **ea3** | **ea3** | **5257** |
| **D1.2** | **D1.1 descendant** | **H5N1 (am-NA)** | **am24** | **ea3** | **am5** | **ea3** | **am13** | **am4N1** | **ea3** | **ea3** | **5** |
| **D1.4** | **D1.1 descendant** | **H5N1 (am-NA)** | **am3.2** | **ea3** | **am4** | **ea3** | **am13** | **am4N1** | **ea3** | **ea3** | **14** |
| **D1.5** | **D1.1 descendant** | **H5N1 (am-NA)** | **am24** | **ea3** | **am4** | **ea3** | **am13** | **am10N1** | **ea3** | **ea3** | **5** |
| **D1.6** | **D1.1 descendant** | **H5N2** | **am2.2** | **am22** | **am4** | **ea3** | **am13** | **am11N2** | **ea3** | **ea3** | **5** |
| **D1.7** | **D1.1 descendant** | **H5N1 (am-NA)** | **am24** | **ea3** | **am12** | **ea3** | **am16** | **am4N1** | **ea3** | **ea3** | **10** |
| **D1.8** | **D1.1 descendant** | **H5N1 (am-NA)** | **am24** | **ea3** | **am4** | **ea3** | **am1.1** | **am4N1** | **ea3** | **ea3** | **5** |
| **D1.9** | **D1.1 descendant** | **H5N1 (am-NA)** | **am2.2** | **ea3** | **am14** | **ea3** | **am13** | **am4N1** | **ea3** | **ea3** | **12** |
| **D1.10** | **D1.1 descendant** | **H5N1 (am-NA)** | **am2.2** | **ea3** | **am12** | **ea3** | **am1.6** | **am4N1** | **ea3** | **ea3** | **2** |
| **D1.12** | **D1.1 descendant** | **H5N1 (am-NA)** | **am18** | **ea3** | **am12** | **ea3** | **am13** | **am4N1** | **ea3** | **ea3** | **3** |
| **D2.1** | **D1.1 descendant** | **H5N2** | **am25** | **ea3** | **am4** | **ea3** | **am1.1** | **am5N2** | **ea3** | **ea3** | **2** |
| Minor101 | D1.1 descendant | H5N9 | am24 | ea3 | am4 | ea3 | am11 | am7N9 | ea3 | ea3 | 1 |
| Minor102 | D1.1 descendant | H5N5 | am1.1 | ea3 | am8 | ea3 | am13 | am8N5 | ea3 | ea3 | 1 |
| Minor107 | D1.1 descendant | H5N2 | am23 | ea3 | am4 | ea3 | am13 | am9N2 | ea3 | ea3 | 1 |
| Minor110 | D1.1 descendant | H5N1 (am-NA) | am24 | ea3 | am8 | ea3 | am15 | am4N1 | ea3 | ea3 | 1 |
| Minor111 | D1.1 descendant | H5N1 (am-NA) | am2.2 | ea3 | am9 | ea3 | am13 | am4N1 | ea3 | ea3 | 12 |
| Minor112 | D1.1 descendant | H5N1 (am-NA) | am24 | am9 | am4 | ea3 | am13 | am4N1 | ea3 | ea3 | 1 |
| Minor114 | D1.1 descendant | H5N1 (am-NA) | am24 | ea3 | am9 | ea3 | am13 | am4N1 | ea3 | ea3 | 9 |
| Minor120 | D1.1 descendant | H5N1 (am-NA) | am2.2 | am25 | am4 | ea3 | am13 | am4N1 | ea3 | am1.2 | 1 |
| Minor124 | D1.1 descendant | H5N1 (am-NA) | am24 | ea3 | am4 | ea3 | am17 | am4N1 | ea3 | ea3 | 1 |
| Minor129 | D1.1 descendant | H5N1 (am-NA) | am18 | ea3 | am8 | ea3 | am9 | am4N1 | ea3 | ea3 | 1 |
| Minor131 | D1.1 descendant | H5N1 (am-NA) | am24 | ea3 | am12 | ea3 | am13 | am4N1 | ea3 | ea3 | 8 |
| Minor133 | D1.1 descendant | H5N1 (am-NA) | am24 | ea3 | am14 | ea3 | am13 | am4N1 | ea3 | ea3 | 1 |
| Minor135 | D1.1 descendant | H5N1 (am-NA) | am18 | ea3 | am4 | ea3 | am13 | am4N1 | ea3 | ea3 | 1 |
| Minor137 | D1.1 descendant | H5N1 (am-NA) | am24 | ea3 | am4 | ea3 | am6 | am4N1 | ea3 | ea3 | 2 |
| Minor16 | A1 descendant | H5N4 | am2.2 | am3 | ea1 | ea1 | am3 | am1N4 | ea1 | ea1 | 1 |
| Minor36 | A1 descendant | H5N5 | am1.3 | am1.3 | am1.2 | ea1 | am1.2 | am2N6 | ea1 | ea1 | 1 |
| Minor97 | D1.1 descendant | H5N1 (am-NA) | am24 | ea3 | am4 | ea3 | ea3 | am4N1 | ea3 | ea3 | 4 |
| Minor98 | A3 descendant | H5N2 | am2.1 | ea3 | am7 | ea3 | am14 | am6N2 | ea3 | ea3 | 6 |
| Minor99 | D1.1 descendant | H5N1 (am-NA) | am2.2 | ea3 | am4 | ea3 | am13 | am4N1 | ea3 | ea3 | 1 |

**Table S2.** Genotype-defining residues among dominant reassortant genotypes. Only residues found in >90% of all sequences within the genotype but <20% of other major genotypes in the table and <10% of all other clade 2.3.4.4b sequences combined are listed. All residues follow sequential numbering except HA, which follows an H5 numbering scheme. White cells represent Eurasian segment lineages while colored cells indicate lineages acquired from North American LPAI, with cells shaded with the same color indicating a shared origin between genotypes.

| Genotype | PB2 | PB1 | PA | HA | NP | NA | MP |  | NS |  |
| --- | --- | --- | --- | --- | --- | --- | --- | --- | --- | --- |
|  |  |  |  |  |  |  | M1 | M2 | NS1 | NEP |
| B3.2 | 340K, 464M | 215K, 429R | -- | -- | -- | -- | -- | -- | -- | -- |
| B3.6 | 251K, 353R, 663R, 667I | -- | -- | -- | 230L | -- | -- | -- | -- | -- |
| B3.13 | 109I, 362G, 441N, 495I, 631L, 649I, 676A | -- | 219I, 497R | -- | 482N | -- | -- | -- | 7L | 7L |
| C2.1 | 129A, 470N | 386K, 618G | 104R, 609R | 226T, 503Y | 353I, 450S | -- | -- | -- | 70G, 111M, 114P, 161T, 176D | -- |
| D1.1 | -- | 16D, 154S, 207R, 614D | -- | 11I, 52A, 120L, 226V, 341R, 491D, 526I | -- | 20I, 23V, 44N, 45H, 48T, 53V, 74L, 75I, 81D, 82P, 84A, 221S, 234I, 241I, 257R, 286S, 287E, 288V, 329S, 336G, 338V, 395A | 200A | 61R | 75G, 147I, 171N, 193Q | 36K, 63E |

**Table S3.** D1.1-specific NA segment residues by protein domain.

| NA domain<br>(residue position<br>range) | Sequential (N1)<br>numbering | N2 numbering |
| --- | --- | --- |
| Tail (1-6) | -- | -- |
| Transmembrane<br>(7-29) | I20 | I20 |
|  | V23 | V23 |
| Stalk (30-84) | N44 | N44 |
|  | H45 | H45 |
|  | T48 | insertion |
|  | V53 | V51 |
|  | L74 | L72 |
|  | I75 | L73 |
|  | D81 | D79 |
|  | P82 | P80 |
|  | A84 | A82 |
| Head (85+) | S221 | S220 |
|  | I234 | I233 |
|  | I241 | I240 |
|  | R257 | R256 |
|  | S286 | S285 |
|  | E287 | E286 |
|  | V288 | V287 |
|  | S329 | S329 |
|  | G336 | G339 |
|  | V338 | V341 |
|  | A395 | A399 |

**Table S4.** Comparison of dispersal metrics estimated for recent H5N1 genotypes having spread in North and South America. We report the weighted diffusion coefficient (WDC) estimated for each of the four dominant H5N1 reassortant genotypes in the Americas (B3.2, B3.6, C2.1, and D1.1). For each dataset, we report both the posterior median estimate and the 95% highest posterior density (HPD) interval. For comparison, we also report the same dispersal metric previously estimated for the West Nile virus spread in North America.

| Dataset | Weighted diffusion coefficient (km<sup>2</sup>/day) | WDC during invasion (km<sup>2</sup>/ day) | WDC after invasion (km<sup>2</sup>/ day) |
| --- | --- | --- | --- |
| HPAI H5N1 (B3.2) | 3030 [2741, 336431] | 3418 [3087, 3850] | 1908 [1358, 2581] |
| HPAI H5N1 (B3.6) | 2878 [2653, 3126] | 3200 [2677, 3884] | 2074 [1385, 2794] |
| HPAI H5N1 (C2.1) | 1659 [1376, 1950] | 1867 [868, 3475] | 536 [296, 815] |
| HPAI H5N1 (D1.1) | 2494 [2268, 2736] | 5873 [4248, 7949] | 1949 [1711, 2218] |
| West Nile virus (North America) | 161 [151, 170] | 312 [280, 373] | 119 [107, 135] |

**Table S5.** Equivalent designations between GenoFLU and new global nomenclature system ggFLU (. https://github.com/YiSong89/ggFlu)

| HA clade | Major genotype (ggFLU) | GenoFlu |
| --- | --- | --- |
| 2.3.4.4b | B.16.1 | A1, A2 |
| 2.3.4.4b | B.16 | A2 |
| 2.3.4.4b | G.4 | A3 |
| 2.3.4.4b | G.6 | A3 |
| 2.3.4.4b | G.6.2 | A3 |
| 2.3.4.4b | G.8 | A3 |
| 2.3.4.4b | B.23.2 | A3, A5 |
| 2.3.4.4b | B.31 | A4 |
| 2.3.4.4b | B.24 | A6 |
| 2.3.4.4b | E.4 | B1.1 |
| 2.3.4.4b | E.6 | B1.2 |
| 2.3.4.4b | E.6 | B1.3 |
| 2.3.4.4b | E.5 | B2.1 |
| 2.3.4.4b | L.1 | B2.2 |
| 2.3.4.4b | E.1 | B3.1 |
| 2.3.4.4b | K.1 | B3.10 |
| 2.3.4.4b | K.1 | B3.11 |
| 2.3.4.4b | K.1 | B3.12 |
| 2.3.4.4b | E.2 | B3.13 |
| 2.3.4.4b | E.2 | B3.2 |
| 2.3.4.4b | E.2 | B3.3 |
| 2.3.4.4b | K.2 | B3.4 |
| 2.3.4.4b | E.2 | B3.5 |
| 2.3.4.4b | K.1 | B3.6 |
| 2.3.4.4b | E.2 | B3.7 |
| 2.3.4.4b | K.1 | B3.8 |
| 2.3.4.4b | E.2 | B3.9 |
| 2.3.4.4b | E.1 | B4.1 |
| 2.3.4.4b | E.3 | B5.1, Minor09 |
| 2.3.4.4b | E.12 | C1.1 |
| 2.3.4.4b | E.13 | C2.1 |
| 2.3.4.4b | E.15 | C3.1 |
| 2.3.4.4b | G.6.5 | D1.1 |
| 2.3.4.4b | P.4 | D1.2 |
| 2.3.4.4b | P.5 | D1.3 |
| 2.3.4.4b | P.6 | D2.1 |

**Figure 1.**
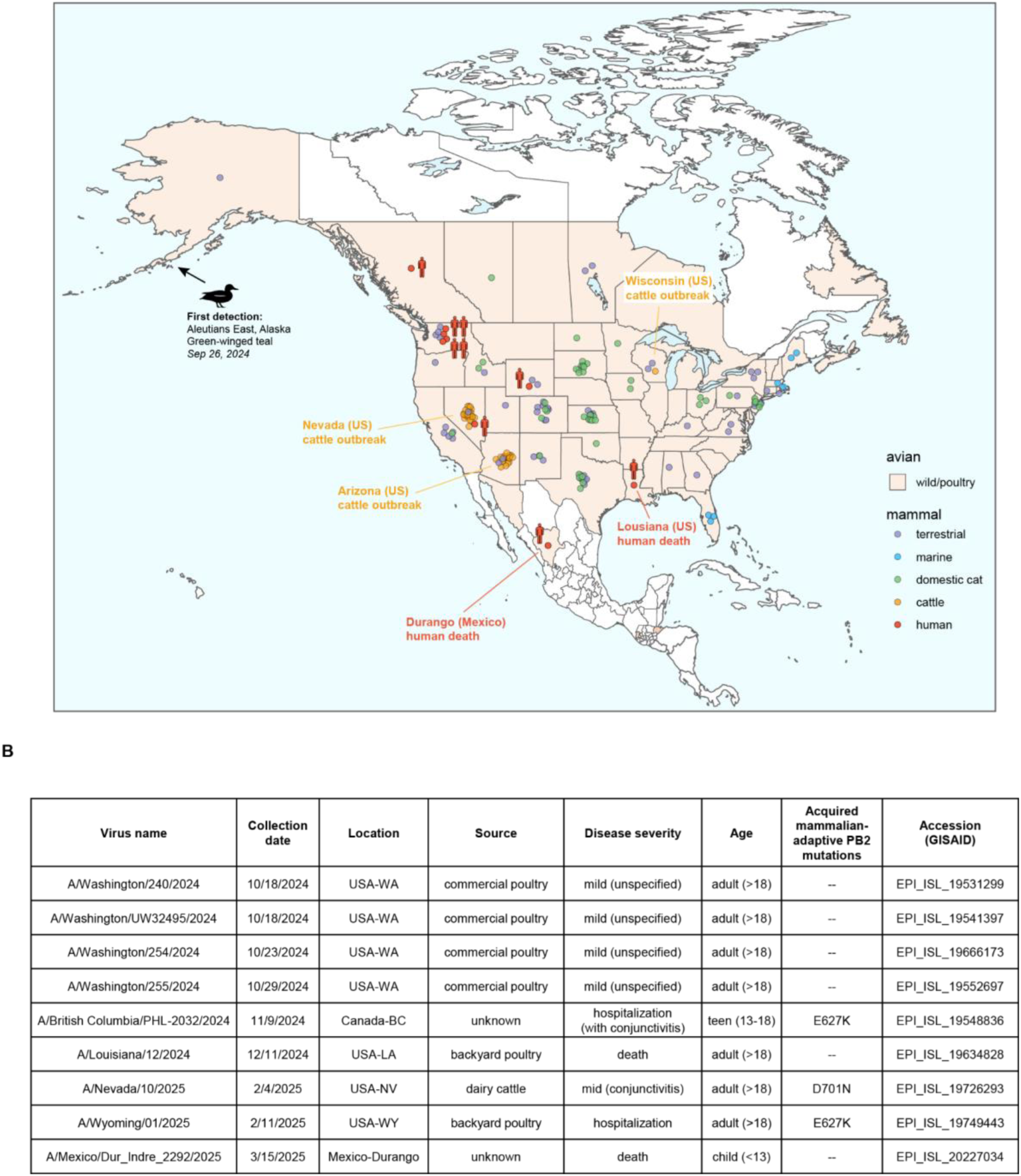
H5N1 genotype D1.1 sample distribution across North America, 2024-2025. (A) Map of North America showing the geographic distribution of publicly available D1.1 samples collected in 2024-2025 across the US, Canada and Mexico. Circles denote mammalian samples and are colored according to the host type. The coordinates for each data point are not exact and are jittered around the centroid for each US state, Canadian province and Mexican state. Level 1 administrative regions from where wild and/or poultry avian samples have been collected are shaded in beige while regions where D1.1 has not been detected are shaded in white. (B) Description of D1.1 infections in humans that were sampled and sequenced.

**Figure S2.**
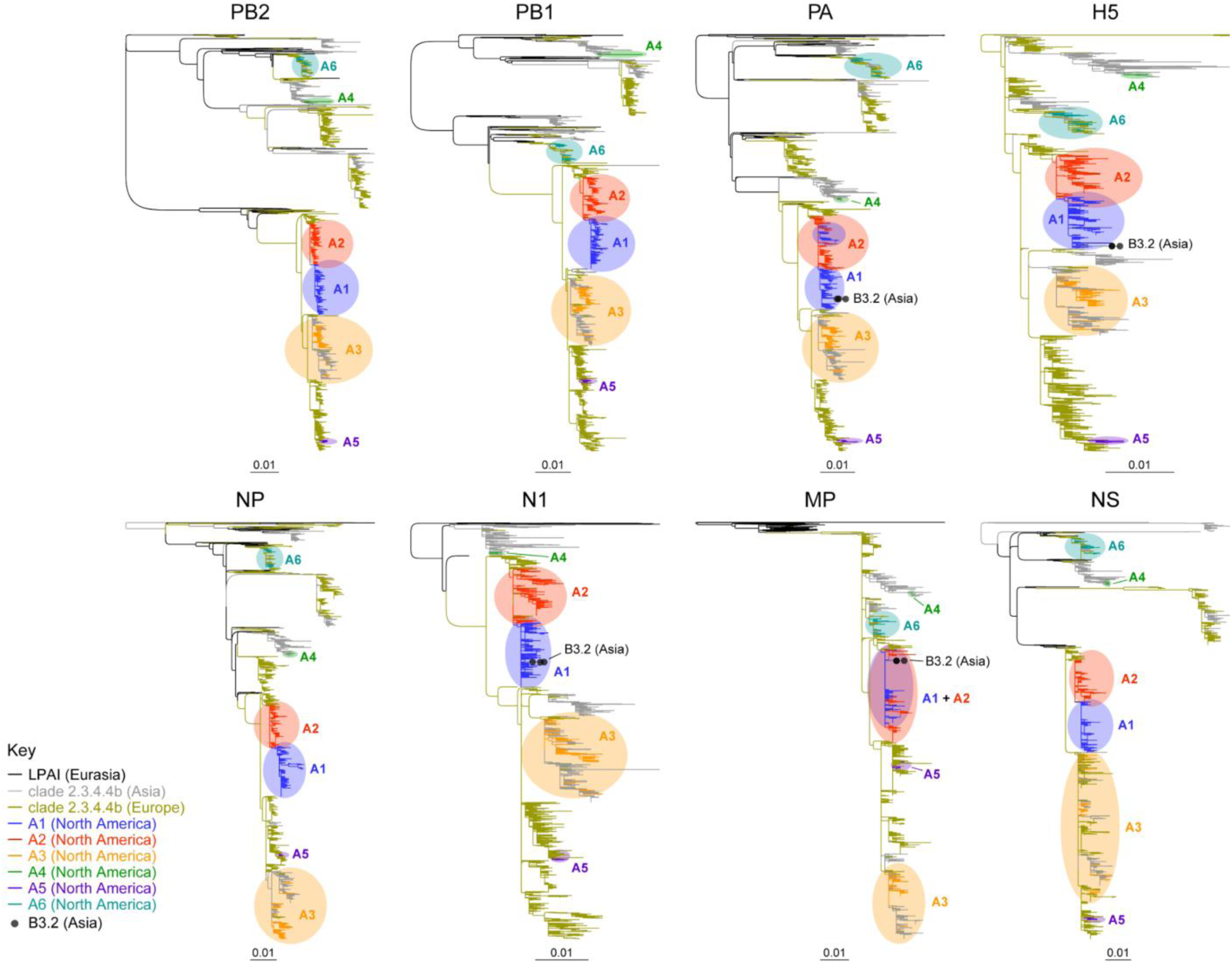
Midpoint-rooted segment phylogenies for EA genotypes (A1-A6) sampled in the Americas, and Eurasian LPAI and clade 2.3.4.4b sequences (collected between January 1, 2015 and June 13, 2025). Tips for North American reassortant genotype B3.2 samples collected in Asia are circled. Full dataset tree files available at https://github.com/nidiatrovao/H5N1Americas.

**Figure S3.**
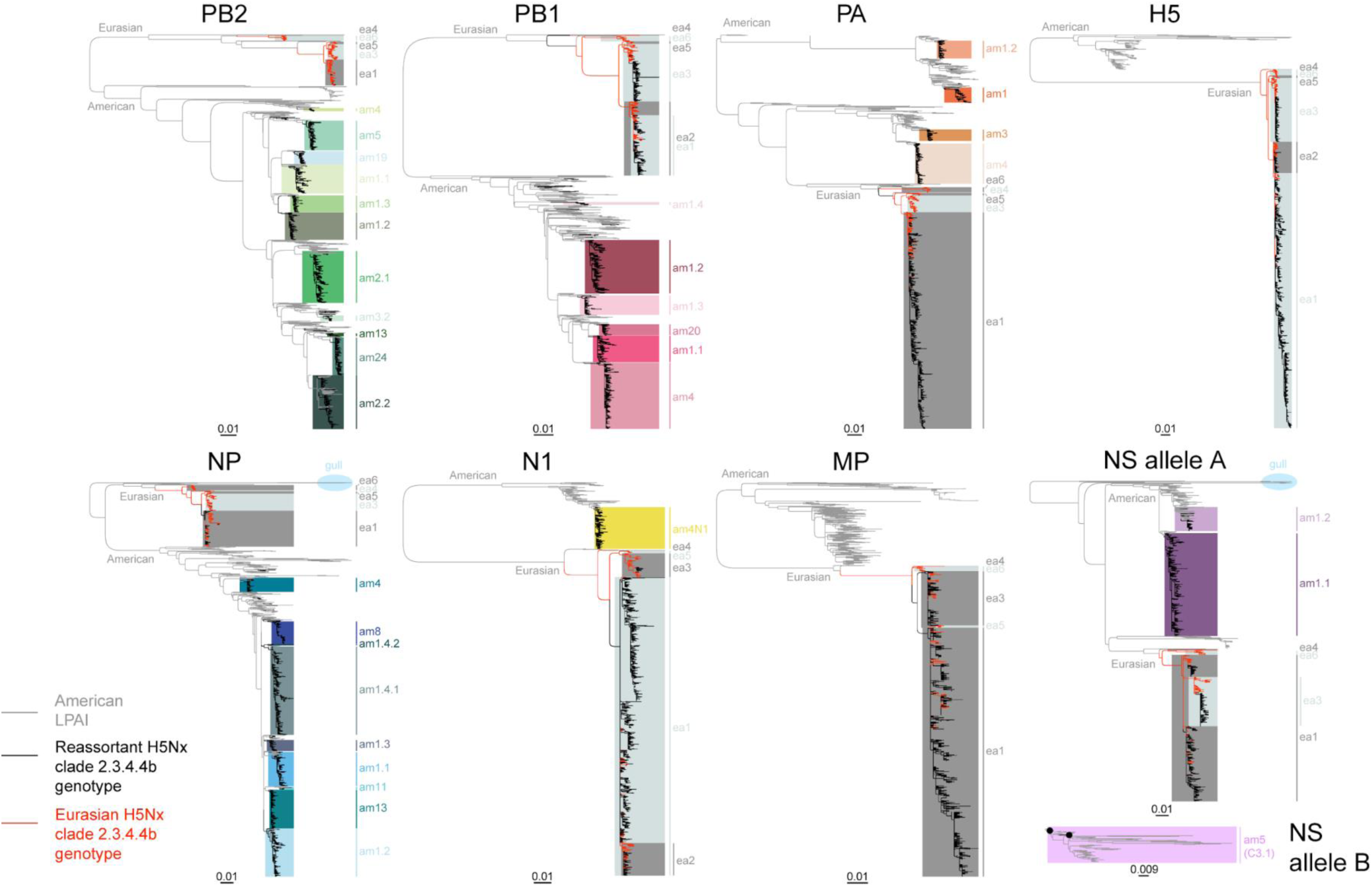
Subsampled, midpoint-rooted segment phylogenies for H5Nx clade 2.3.4.4b and LPAI viruses sampled in the Western hemisphere (collected between January 1, 2015 and June 13, 2025). Full and subsampled dataset tree files are available at https://github.com/nidiatrovao/H5N1Americas

**Figure S4.**
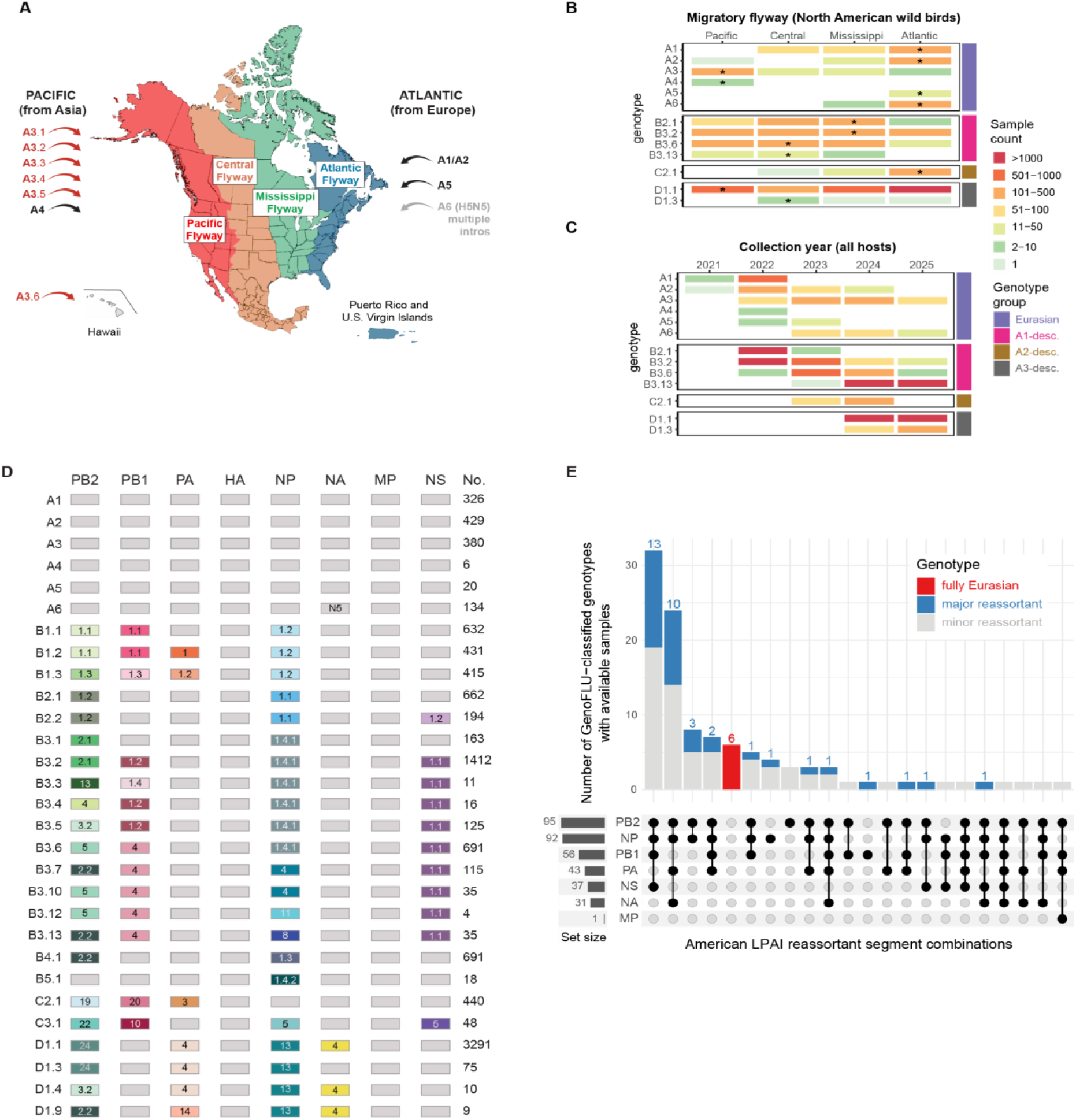
(A) Map of North America’s four administrative migratory flyways, defined by the US Fish and Wildlife Service. Arrows indicate independent introductions of Eurasian H5Nx clade 2.3.4.4b genotypes into North American migratory flyways, where black arrows indicate single H5N1 introductions, red arrows indicate repeated introductions of genotype A3, and the grey arrow indicates multiple introductions of H5N5 genotype A6. (B) Heatmap shaded by the number of sequences in each North American migratory flyway for North American wild birds (*n* = 7,869) and (C) heatmap shaded by collection year across all hosts (*n* = 17,243). Only select genotypes are shown. Asterisks denote the flyway where the initial detection occurred. (D) Genotype constellation for clade 2.3.4.4b H5N1 major genotypes with >10 sequences available. Grey squares indicate Eurasian lineage; colored squares represent reassorted North American LPAI lineages. The number of genome sequences available from wild birds in the Americas is provided for each genotype in the last column. (E) The y-axis represents the number of H5Nx genotypes with available sequences (defined by GenoFlu; *n* = 107) with each combination of LPAI segments. Major reassortant genotypes (as defined by GenoFlu, see Methods) are shaded blue; minor reassortant genotypes are shaded grey; non-reassorted fully Eurasian genotypes are shaded red. The count of genotypes with an LPAI-derived segment for each of the eight gene segments is indicated on the left.

**Figure S5.**
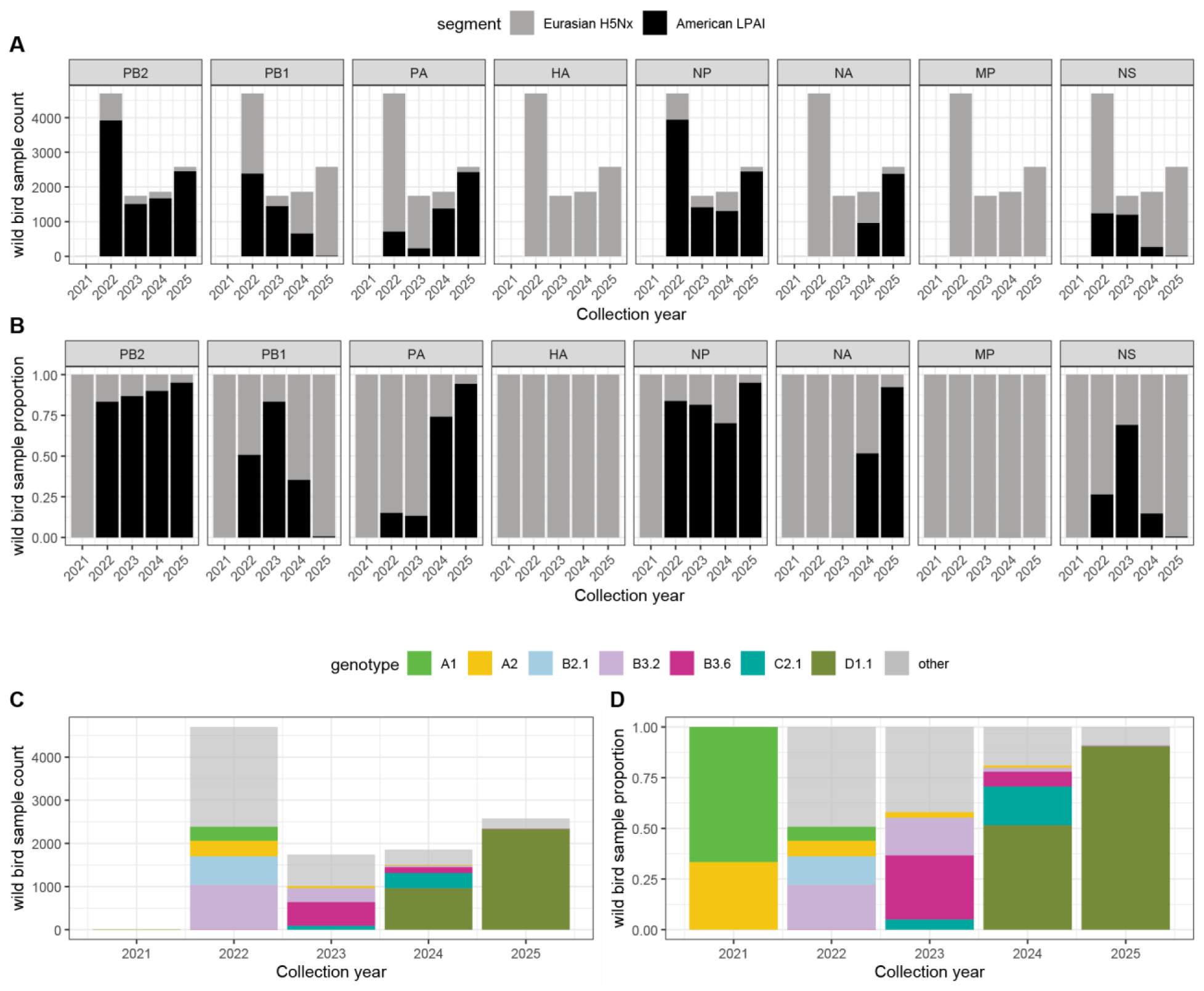
Yearly counts and proportions for H5Nx clade 2.3.4.4b segment sequences during 2021-2025. (A) Yearly counts and (B) proportions for H5Nx clade 2.3.4.4b wild bird segment sequences shaded by Eurasian H5Nx (blue) or American LPAI (black) origin. **(C)** Yearly counts and (D) proportions for H5Nx clade 2.3.4.4b wild bird samples shaded by genotype.

**Figure S6.**
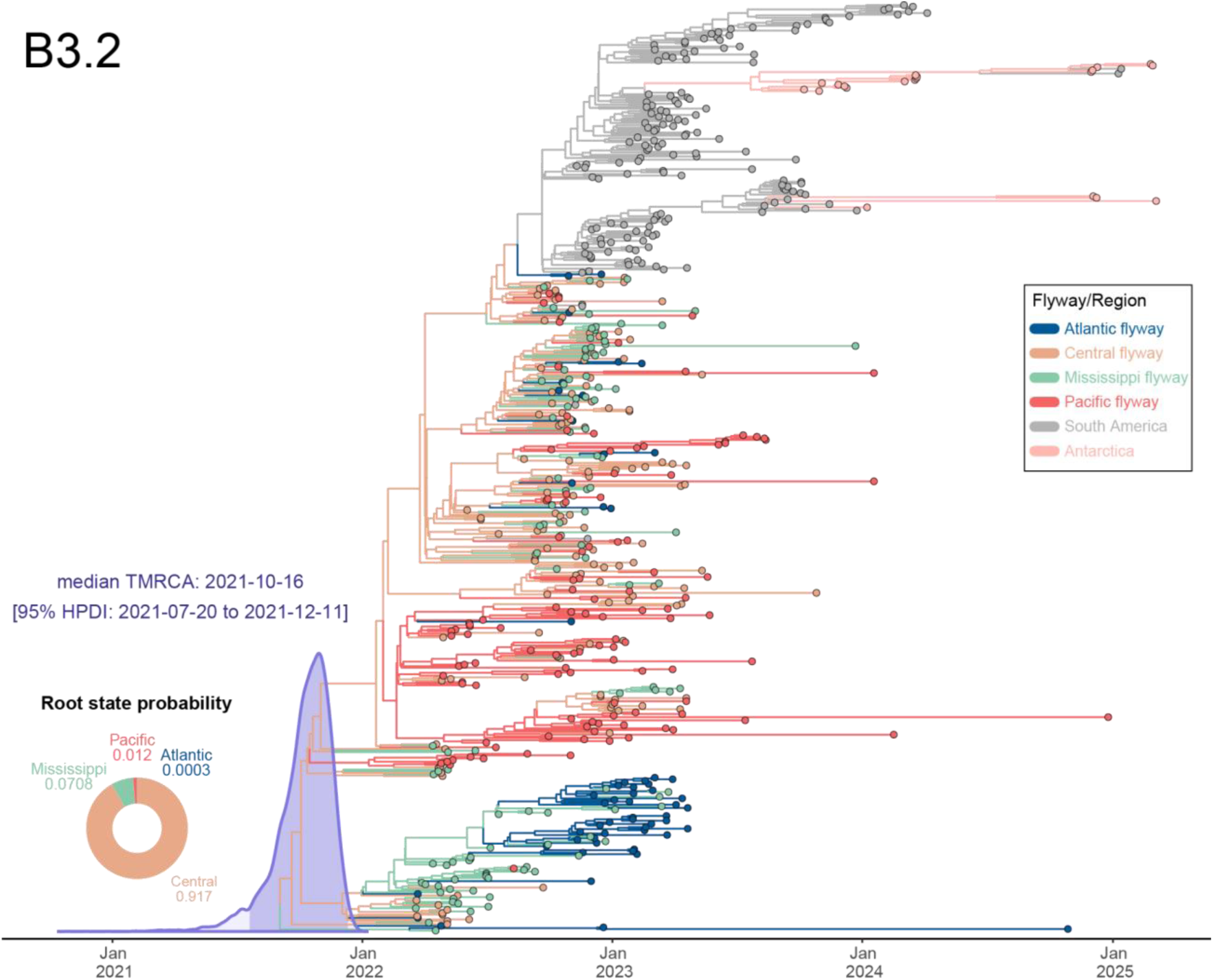
Discrete phylogeographic reconstruction of the dispersal history for H5N1 genotype B3.2 across the North American waterfowl migratory flyways. The consensus phylogenetic tree shows the evolutionary relationships of genotype B3.2 viruses, with the X-axis scaled in years. Tips are shaded according to the region or North American flyway where they were collected, and branches are shaded according to the flyway inferred at ancestral nodes. The purple curve at the root of the tree represents the posterior density distribution for the TMRCA. The donut plot at the base of the phylogeny indicates the posterior flyway root state probabilities.

**Figure S7.**
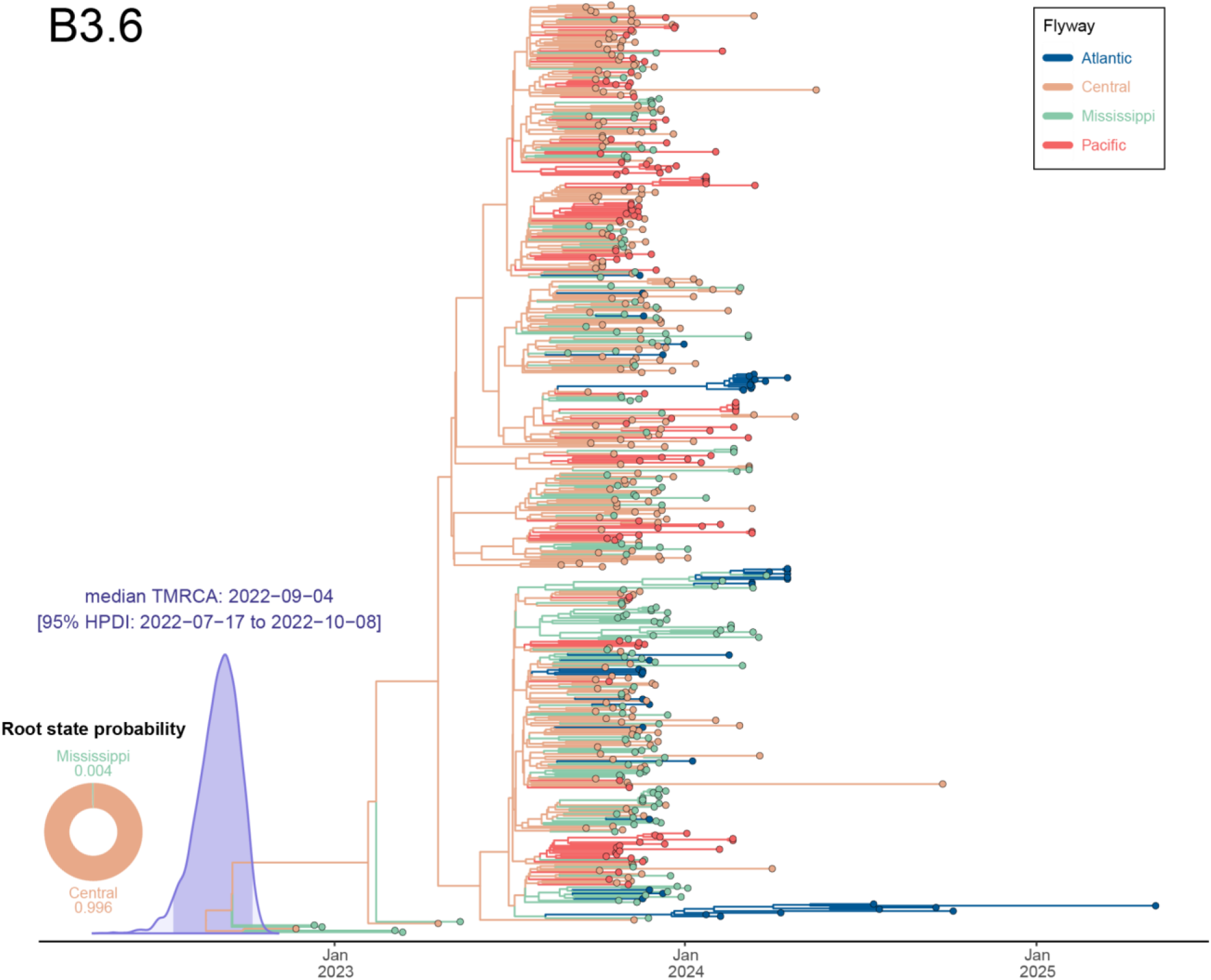
Discrete phylogeographic reconstruction of the dispersal history for H5N1 genotype B3.6 across the North American waterfowl migratory flyways. The consensus phylogenetic tree shows the evolutionary relationships of genotype B3.6 viruses, with the X-axis scaled in years. Tips are shaded according to the flyway where they were collected, and branches are shaded according to the flyway inferred at ancestral nodes. The purple curve at the root of the tree represents the posterior density distribution for the TMRCA. The donut plot at the base of the phylogeny indicates the posterior flyway root state probabilities.

**Figure S8.**
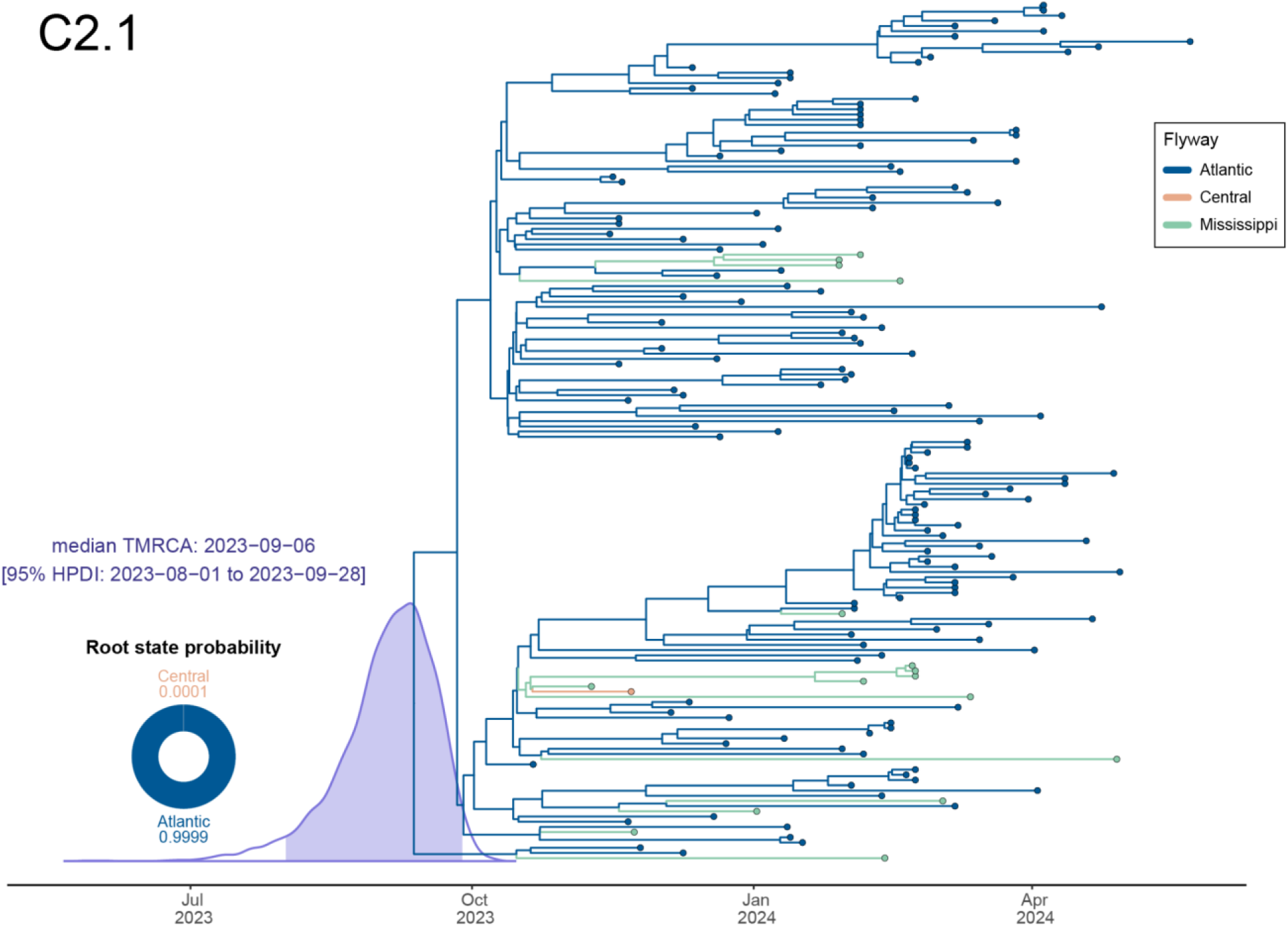
Discrete phylogeographic reconstruction of the dispersal history for H5N1 genotype C2.1 across the North American waterfowl migratory flyways. The consensus phylogenetic tree shows the evolutionary relationships of genotype C2.1 viruses, with the X-axis scaled in years. Tips are shaded according to the flyway where they were collected, and branches are shaded according to the flyway inferred at ancestral nodes. The purple curve at the root of the tree represents the posterior density distribution for the TMRCA.

**Figure S9.**
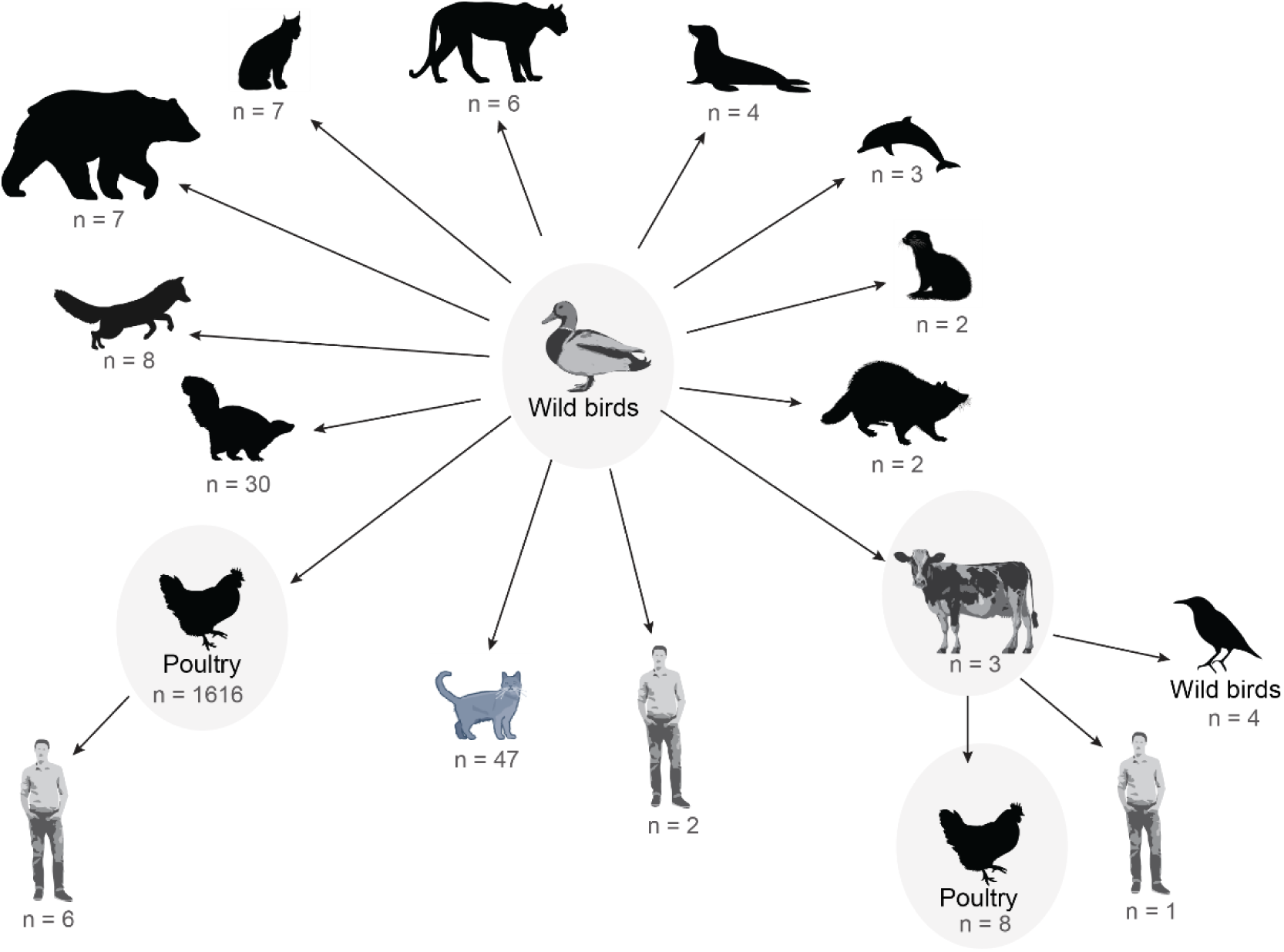
D1.1 transmission routes. Sample sizes represent the number of sequences in each host category.

**Figure S10.**
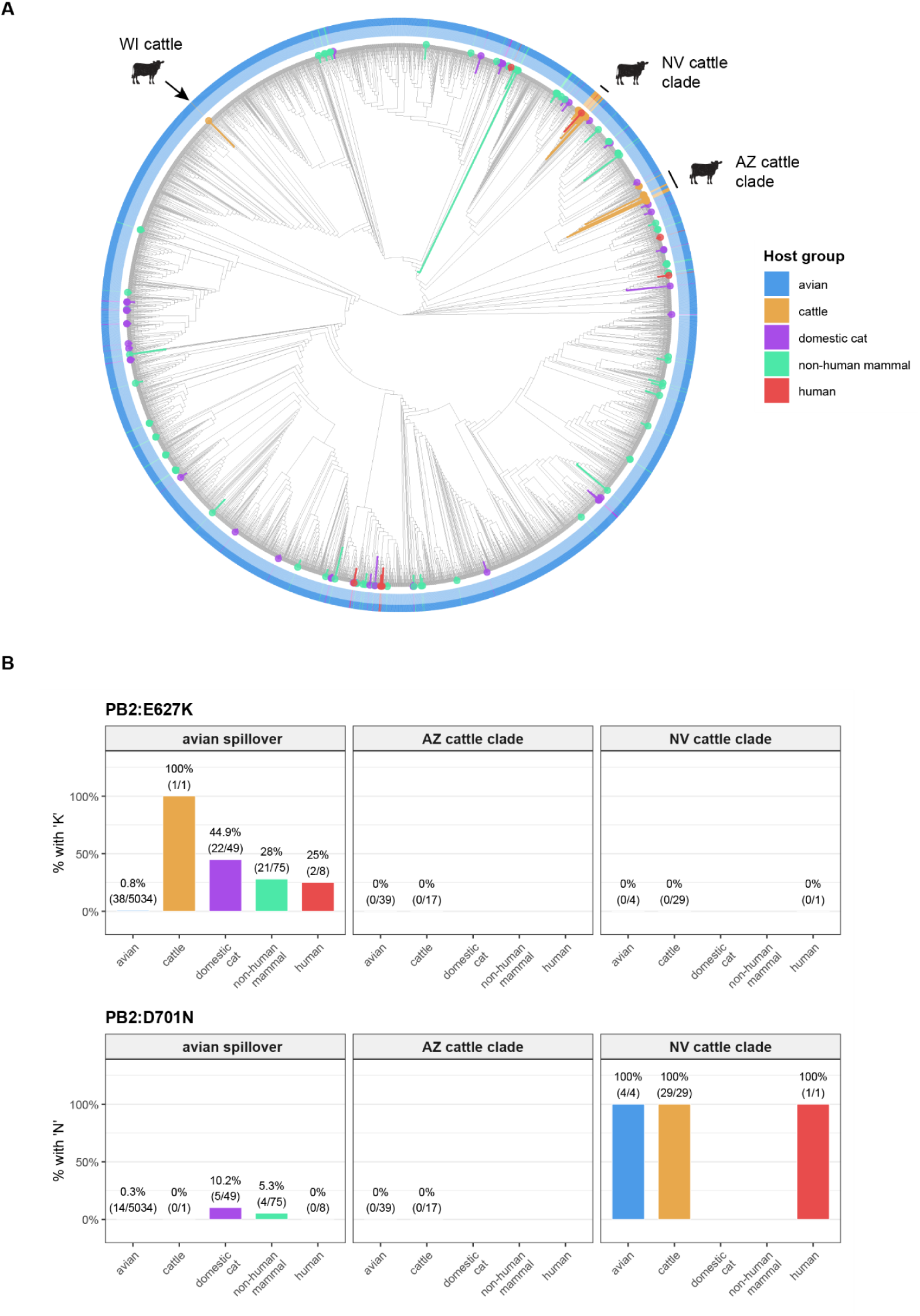
D1.1 spillover. (A) Midpoint-rooted D1.1 ML phylogeny with host group annotations (*n* = 5,257). All three cattle outbreaks are highlighted. (B) Presence of PB2 mammalian-adaptive substitutions E627K and D701N in sequences obtained from different host groups and facetted by transmission route: from cattle (i.e., within “AZ cattle clade” and “NV cattle clade”), and from avian hosts (“avian spillover”). Full PB2 FluMut outputs available at: https://github.com/nidiatrovao/H5N1Americas.

**Figure S11.**
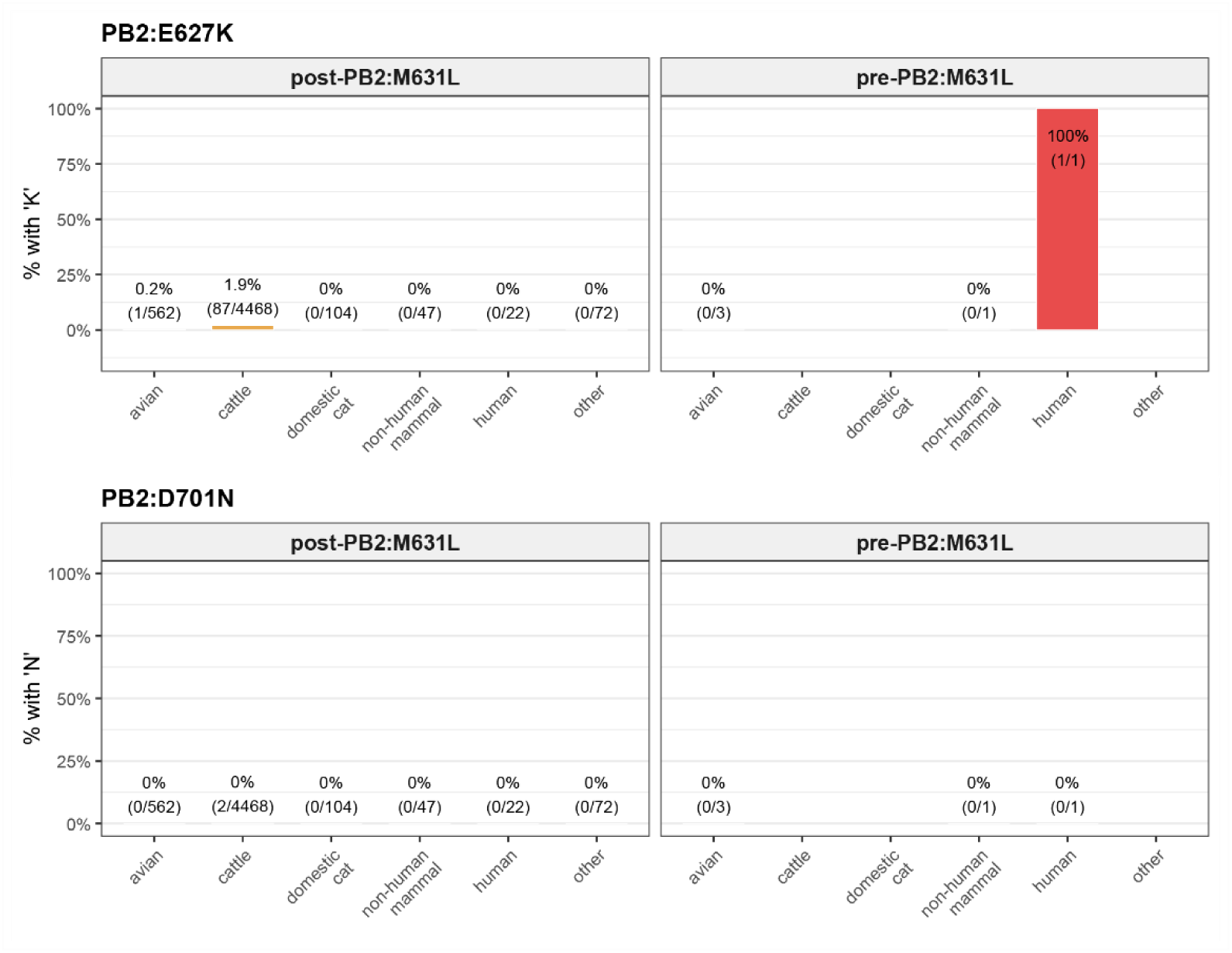
Low frequency of PB2 mammalian-adaptive substitutions E627K and D701N among B3.13 samples. Presence of PB2 mammalian-adaptive substitutions E627K and D701N in sequences obtained from different host groups. Full PB2 FluMut outputs available at: https://github.com/nidiatrovao/H5N1Americas.

